# Algorithm-Powered Analyzer for Continuous Electrochemistry: A Toolkit for Real-Time Electrochemical Data Analysis

**DOI:** 10.1101/2025.09.28.678418

**Authors:** Yinke Jiang, Yihang Chen, Yufei Cai, Keyang Zhou, Hayden J. Ousley, Jun Li, H. Tom Soh, Kaiyu X. Fu

## Abstract

Real-time biosensors offer significant potential for continuous monitoring of biomolecules. However, their practical application and further development face challenges on data analysis, including poor signal-to-noise ratio when effective sensing area decreases due to signal degradation by biofouling, time-consuming and subjective process of manual or semi-automated peak identification, and inconsistencies in data interpretation, thereby complicating reproducibility and cross-comparison of biosensing results. In this study, we introduce the Algorithm-Powered Analyzer for Continuous Electrochemistry (A-PACE), an open-source toolkit providing streamlined and automated data analysis protocol optimized for real-time electrochemical data analysis. A-PACE comprises three modules: (1) Change point detection for automated identification of peak regions; (2) Baseline fitting with multiple algorithms to handle diverse electrochemical signals and baseline screening to eliminate unreasonable fits; (3) Large dataset input with user-friendly interface for data processing, exporting and visualization. To ensure optimal performance, we curated a training set of 2000 electrochemical curves from an extensive electrochemical dataset (>100,000 curves) collected over the past five years under varied electrochemical measurement conditions. These curves were analyzed using 2046 algorithm sets to identify a default algorithm set, compared to existing tools, demonstrating its capability in peak detection and baseline fitting across a broad spectrum of electrochemical data. Case studies reveal the A-PACE can analyze month-long *in vitro* serum data and week-long *in vivo* intravenous blood data, extending the operational lifespan of real-time sensors. This cross-platform compatible toolkit supports both real-time and post-processing analysis, reducing the processing time from minute level per signal by human labeling to second level by A-PACE and subjectivity associated with electrochemical signal processing. By providing this solution for continuous electrochemical data analysis, A-PACE enhances biosensors’ applications in medical diagnostics and continuous monitoring with high throughput analysis.

## INTRODUCTION

The world’s aging population and rising healthcare costs are accelerating a shift toward affordable, decentralized healthcare^1^. Continuous health monitoring enables earlier interventions and better chronic-disease management^2,3^, making real-time biosensors central to this vision. Electrochemical sensors are particularly attractive because they are inexpensive, easily miniaturized, rapid response, and compatible with multiplexed, wearable or implantable formats^29–31^. Continuous electrochemistry inherently generates time-series redox data. Direct techniques such as chronoamperometry (CA)—the basis of continuous glucose monitor (CGM)—relate current to analyte concentration in real time but lack redox specificity. Electrochemical aptamer-based (EAB) sensors, which have been validated in molecular monitoring^11–14^, address specificity by tethering a redox tag (commonly methylene blue) to a surface-immobilized aptamer^15,16^. Target binding induces conformational and dynamical changes that modulate electron-transfer kinetics and/or the tag-electrode distance, producing a current change that is specific to the molecular target and sensitive to electrode dimension and geometry. Current-potential curves in voltammetry fall broadly into non-peak (steady state) curves and peak-shaped (transient) classes^32^. Cyclic voltammetry (CV), differential pulse voltammetry (DPV) and square wave voltammetry (SWV)—workhorses of electrochemical sensing—primarily yield peak-shaped redox currents. In EAB measurements, SWV is most common: an alternating potential drives the reversible redox event of the tethered tag, producing a single, well-defined peak. Target binding alters the aptamer’s conformation and electron transfer rate, shifting the peak’s amplitude and sometimes its position—an ideal handle for analyte quantification.^33^ Accurate analysis therefore requires reliable peak localization and reasonable baseline estimation so that peak height, width and position can be extracted via baseline subtration^33^. In practice, the SWV peak shape and magnitude vary substantially with waveform parameters, electrode dimensions, and matrix conditions (e.g., protein content, fouling, ionic strength)^34,35^. This variability renders fixed-form baseline fits brittle and lab-dependent. What is needed is a flexible yet standardized, objective, and automated approach to peak finding and baseline fitting that is robust to drift and matrix effects, and suitable for both real-time streaming and post-hoc analysis. Continuous *in vivo* sensing faces four persistent challenges despite advances in transduction and acquisition. First, the sensor size-signal tradeoff: miniaturized, wearable/implantable formats generate smaller currents and thus poorer signal-to-noise ratio (SNR)^4–9^. Second, progressive biofouling: unlike end-point measurement, continuous measurements suffer from non-specific adsorption, enzymatic degradation, and foreign-body reaction that stealthily distort signals over time^10^. Third, labor-intensive workflow: long experiments yield massive datasets that demand manual sorting, grouping, and annotation. Fourth, subjective peak detection: examiner-defined settings lead to inconsistent peak-region and peak-shape across software packages, a problem amplified for tiny (challenge 1) and degrading (challenge 2) signals that dominate continuous monitoring, where small misclassifications cascade through subsequent analysis (challenge 3). Hardware and material progress help but do not eliminate these issues: nanostructured electrodes provide sub-millimeter footprints with enhanced currents^17–19^. Native aptamers remain vulnerable to nuclease attack^20^, causing drift in the biofluidic environment. Surface engineering—including hydrogel barrier^21^, zwitterionic monolayers, DNase/RNase-resistant xenonucleic acids^22^—attenuate fouling and improve SNR. And we have recently extended EAB sensor’s lifetime to one month *in vitro* and one week *in vivo* by seamlessly coating nanoporous gold with a hyperbranched polymer^23^.

However, analysis has become the primary bottleneck to turning these signals into reliable measurements. Widely used scientific graphing tools (OriginPro, GraphPad) and electrochemical suites (MultiTrace, EC-Lab, NOVA) offer manual or semi-automated smoothing, baseline fits, and peak finding. MATLAB toolboxes/Python libraries^24,25^ and real-time signal analysis tools like SACMES^26,27^ add flexibility for electrochemical data. Yet none generalizes to the tiny, drift-prone currents that limit long-term *in vivo* sensing. They rely on fixed shapes or fixed-form baselines, require extensive hand-tuning, and yield lab-dependent outcomes on large streaming datasets. In short, the field lacks a flexible yet standardized, objective, and automated approach to peak finding and baseline modeling that is robust to drift and matrix effects and scalable from real-time operation to post-hoc analysis. This gap motivates a rapid, powerful and reliable tool for biosensor development.

To address these unmet analysis needs in continuous electrochemistry, we present the Algorithm-Powered Analyzer for Continuous Electrochemistry (A-PACE), a toolkit that standardizes peak quantification from time-series redox data and specifically engineered for continuous electrochemical signal analysis. The central innovation of this study is a generalized, reliable pipeline capable of continuously isolating peak heights from electrochemical baselines across large datasets. In detail, we use change-point detection to delineate peak and baseline regions objectively in drifting, low-SNR data. Then we fit a screened ensemble of baseline models rather than a single fixed form, avoiding model misspecification across instruments and matrices. Thereafter, we perform efficient-frontier selection to balance accuracy and temporal stability, yielding reproducible peaks with uncertainty metrics suitable for real-time decision-making and post-hoc analysis. Finally, a sliding-window streaming stabilizer further regularizes live readouts, and a standardized I/O plus GUI makes the workflow portable across labs. All signals are processed with the optimized algorithm set in A-PACE using cost- and time-efficient computing resources by implementing in Python-based algorithms^24^ with a Java-based GUI^28^, leading to an open-source platform that can run on multiple operating systems, including Windows, macOS, and Linux. A-PACE formalizes segmentation, baseline modeling, and model selection as an integrated, verifiable process that is robust to tiny, drift-prone currents that dominate long-term in-vivo sensing. By delivering feedback within seconds, A-PACE delivers a rigorously benchmarked analysis infrastructure that accelerates experiment iteration, enhances device performance, and improves the portability of existing continuous health-monitoring technologies, providing a faster, more reliable, and cost-effective solution for biosensor development.

## RESULTS

To achieve the continuous electrochemical signal analysis with algorithm-powered analyzer (**Figure 1a**), A-PACE implements a unified workflow for peak-shaped signals (**Figure 1b**). Three concentric layers, progressing from core to shell, automate peak extraction during continuous electrochemistry. Layer 1 - Change Point Detection (CPD): partitions each curve into peak and baseline zones, providing accurate baseline subsets for fitting and extract peak information after fitting. Layer 2- Algorithm-Powered Analyzer (A-PA): exhaustively screens algorithm combinations to identify the optimal baseline fitting and robustly extract peak currents. Layer 3 - Data I/O control and graphical user interface (GUI) design: enabling parameter tuning, bulk import of raw data, and export of well-structured results for downstream analysis through a customizable interface. Then such architecture is optimized for speed: a multi-processing engine processes each trace with sub-second speed, enabling month-long *in vitro* human serum and week-long *in vivo* intravenous blood datasets to be processed rapidly and supporting real-time visualization with less than a second delay (**Figure 1c**). As a proof of concept, we focus on continuous electrochemical data produced by EAB sensors.

**Figure 1.**
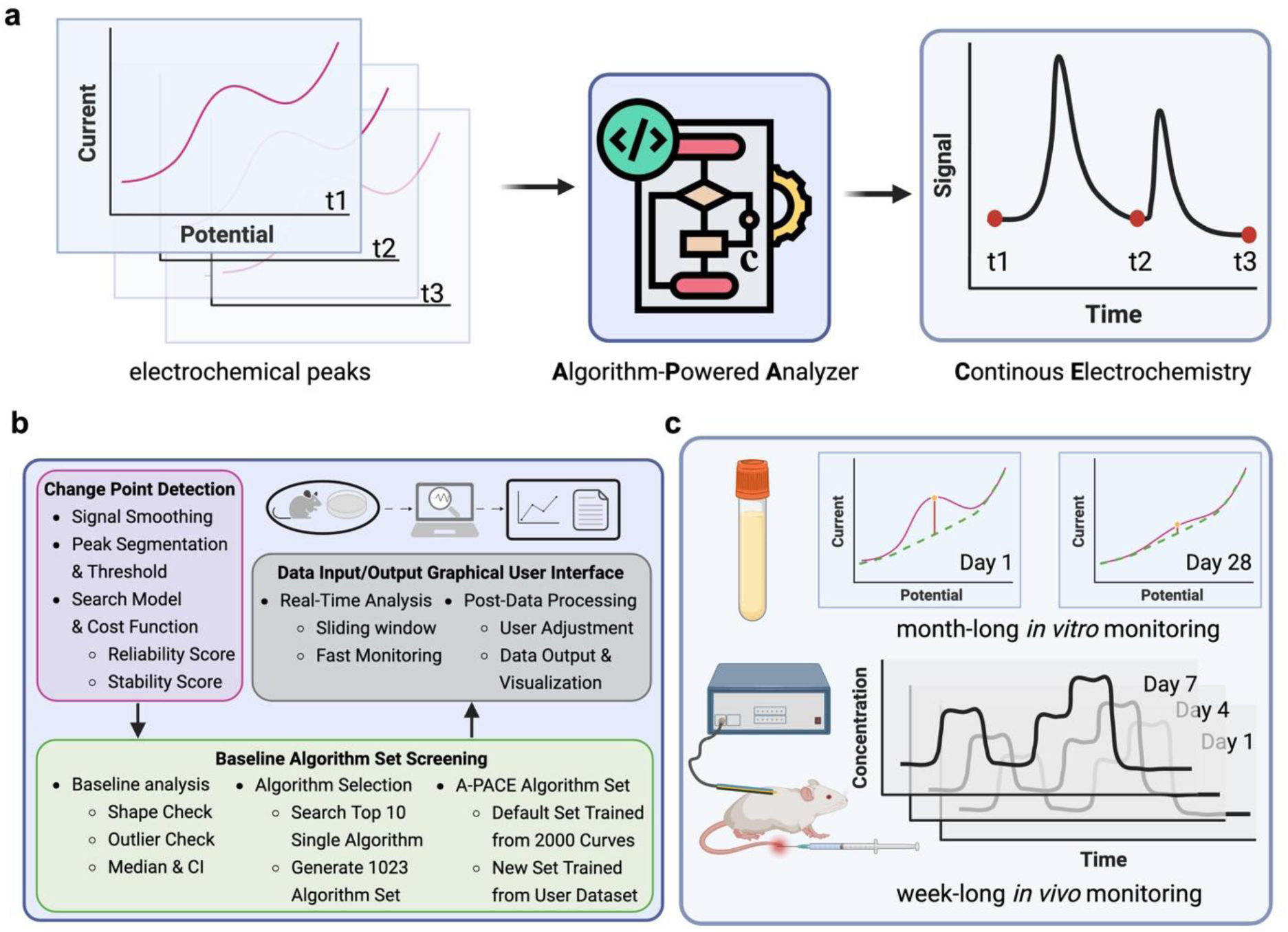
A-PACE pipeline for continuous electrochemical data analysis. **a** Use multiple CPU cores for data processing when multiple files are imported. Generate time-series data based on the time information from files. **b** The data processing workflow of A-PACE, including Layer 1 of Change Point Detection (CPD), providing baseline subsets for fitting, Layer 2 of Algorithm-Powered Analyzer,, screening and selecting algorithm combinations for optimal algorithm set, and Layer 3 of Data I/O control and graphical user interface (GUI) for parameter tuning, raw data import, and final result output. **c** Electrochemical sensors with A-PACE can automatically analyze month-long *in vitro* human serum and week-long *in vivo* intravenous blood datasets.

### Change point detection

Change points (CPs) are specific time points where abrupt increases or decreases happen in a time series, representing transitions between states observed in the experiments. By comparing whether the statistical properties of the “historical segment” and the “current segment” differ significantly, CPD identifies abrupt transitions in time series data^16^. Since early binary-segmentation work in the 1950s^17,18^, CPD has progressed to sophisticated statistical and machine learning algorithms^19–23^ and now underpins applications in finance^24^, traffic analysis^25^, and biosensing^26^. Its maturity and breadth make CPD well-suited for automated SWV peak localization. After importing a SWV data file, A-PACE extracts data information, for example, potential range, current level, time information, and other metadata (channel, frequency, amplitude, step size, etc.). Because SNRs and peak widths differ across instruments and matrices, two user-set parameters guide CPD (**Figure 2a**): (1) noise level and (2) peak region threshold (the maximum fraction of a curve assignable to the peak, default value set as 65%) (more details in the **Supplementary Note 1**). With these two parameters, A-PACE locates two CPs, partitioning each curve into a peak region (between CPs) and a baseline region (outside CPs), as shown in **Figure 2b**. Baseline data are used for baseline fitting and peak data are used for baseline screening and peak height extraction, respectively. For typical SNRs (10-40), level 2 smoothing removes noise without attenuating the peak; level 1 preserves more detail but keeps noise, whereas level 3 over-smooths (**Figure S1**). The peak-region threshold strongly influences CP placement. A 65% threshold can capture 46% of the curve, nearly bracketing the peak; but at 70%, the right CP drifted, incorporating baseline points, and impairing subsequent fitting (**Figure 2c**). To curb artifacts, the minimum CP separation was set to 20% of the curve length (user-adjustable), and the threshold capped peak size.

**Figure 2.**
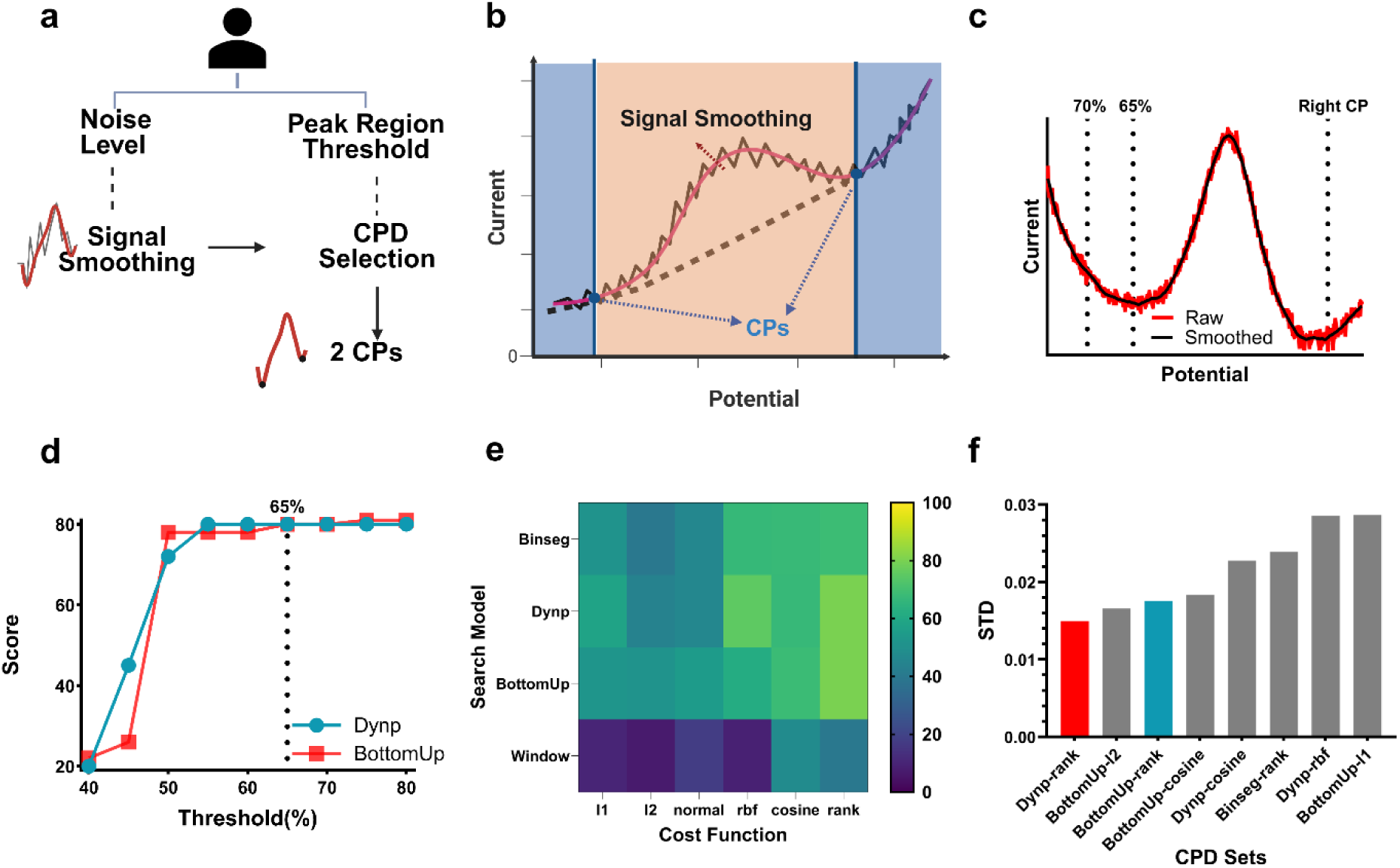
CPD workflow that includes peak region recognition and the the comparison of different CPD algorithms. **a** Two parameters can be set by the user: the noise level and the peak region threshold. These values can affect the selection of the two change points using various CPD algorithms. **b** Savitzky-Golay filter is used for signal smoothing with the following peak region segmentation process defined by two change points. **c** Example of two peak region thresholds (70% vs. 65%) that result in different change points. **d** The performance of the two highest-scoring CPD algorithms as a function of threshold. **e** Scores of four algorithms combined with six cost functions at a threshold of 65%, with Dynp-rank and BottomUp-rank as the top two. **f** The CPD algorithm combinations are ranked by the standard devitation of the change points obtained at a threshold of 65%.

Since the location of CPs directly determines the data used for baseline fitting, it is essential to reliably identify appropriate CPs across signal curves with highly diverse shapes. To achieve this, we systematically evaluated various CPD algorithms on a large set of signals and compared their outputs against manually annotated references. First, we selected 40 folders from a large electrochemical dataset collected over years (details in **Methods**), and each folder includes 50 time-series SWV files. Their SWV signals were collected with different amplitude, frequency and step size with electrode size ranging from single-digit micrometer to sub-millimeter in either static holder or flow cell. The resulted currents span five orders of current magnitude—from single-digit nanoampere to sub-milliampere— with different noise levels (more dataset details in **Figure S2**). Then reliable peak analysis begins with optimal CPD choices: the algorithm and its operating settings determine which samples are treated as baseline versus peak, directly impacting baseline fitting and peak current readout.

Based on our representative dataset, we then bench-marked CPD algorithms from Python-based *Ruptures* package^47^. Four search methods—Dynamic programming (Dynp), Binary segmentation (Binseq), Window sliding segmentation (Window) and Bottom-up segmentation (BottomUp) with six cost functions of each: CostL1 (l1), CostL2 (l2), CostNormal (normal), CostRank (rank), CostCosine (cosine) and CostRbf (rbf), leading to 24 combinations that were applied to 2,000 SWV files for automated CPs extraction, which were compared with human-labeled CPs (details in **Table S1**). The score (from 0 to 1) equaled the percentage of curves whose CPs lay within the annotated region (more details in the **Supplementary Note 2**). Thus, higher scores indicate greater concordance between CPD algorithm-identified CPs and the human-labeled regions. Then, nine thresholds from 40% to 80% (with step size of 5%) with a fixed level 2 smoothing were tested. By sweeping and selecting the optimal threshold of each CPD algorithms, we can reduce algorithm-specific sensitivity to the threshold and can detect whether CPs are too close or too far apart, given that most CP separations lie within 65%. Among all the results (**Figure S3**), Dynp-rank and BottomUp-rank consistently scored highest, plateauing beyond a 65% threshold (**Figure 2d**) and outperforming all other pairs at 65% (**Figure 2e**). Compared with algorithms performance with 60% threshold, the overall scores with 65% threshold increase significantly, which indicates a better fit with human-labeled CPs. However, when the threshold increased to 70%, there are slight differences between many algorithms. To avoid excessively broad peak region, the threshold was set as 65% (more details in **Figure S3**). To assess inter-folder consistency, we computed the standard deviation (STD) of CPs locations; Dynp-rank and BottomUp-rank again exhibited the lowest STDs (**Figure 2f**), confirming superior stability. Therefore, these two methods were selected for subsequent analysis.

### Baseline fitting and screening

With two reliable CPs and sufficient baseline-region identified in the CPD step, baseline fitting was subsequently conducted using a variety of algorithms. Usually, the baseline is determined by single algorithm. And one of the most common algorithms is polynomial fitting, including linear fitting when the order is 0. However, even with polynomial orders up to 15, accuracy remains limited^8^. Therefore, using an algorithm set composed of multiple fitting strategies—such as polynomial, B-spline, and Gaussian fitting—instead of single algorithm enables more robust performance considering the diversity of different baseline shapes, magnitudes and noise levels (more examples in **Figure S4**). We established a unified workflow with multiple steps of screening and filtering processes to select the appropriate and reasonably fitted baselines (**Figure 3a**). First, the peak height is a positive value in our electrochemical setup regardless of reduction or oxidation peaks. Any fitted baseline for which more than 10% of its data exceeded the raw signal in the peak region is discarded. This step excluded fitted curves that do not conform to the baseline definition, thereby precluding any negative peak values (**Figure 3b**). Second, when the detected CPs are narrower than the actual CPs, this leads to a portion of peak-region data points being wrongly included in the baseline fitting. As a result, the concavity of the fitted baseline would rise greatly against our initial assumptions. To mitigate this, the areas under the raw signal curve (AUCs) were calculated, with the straight line connecting the two CPs and the fitted baseline as bottom, respectively. The AUC from the fitted baselines that shown showing abnormal difference (>30%) with the AUC from straight line were excluded (**Figure 3c**) However, some cases shown in **Figure S4**, like panel c, if the AUC from the line connecting the two CPs is negative value,

**Figure 3.**
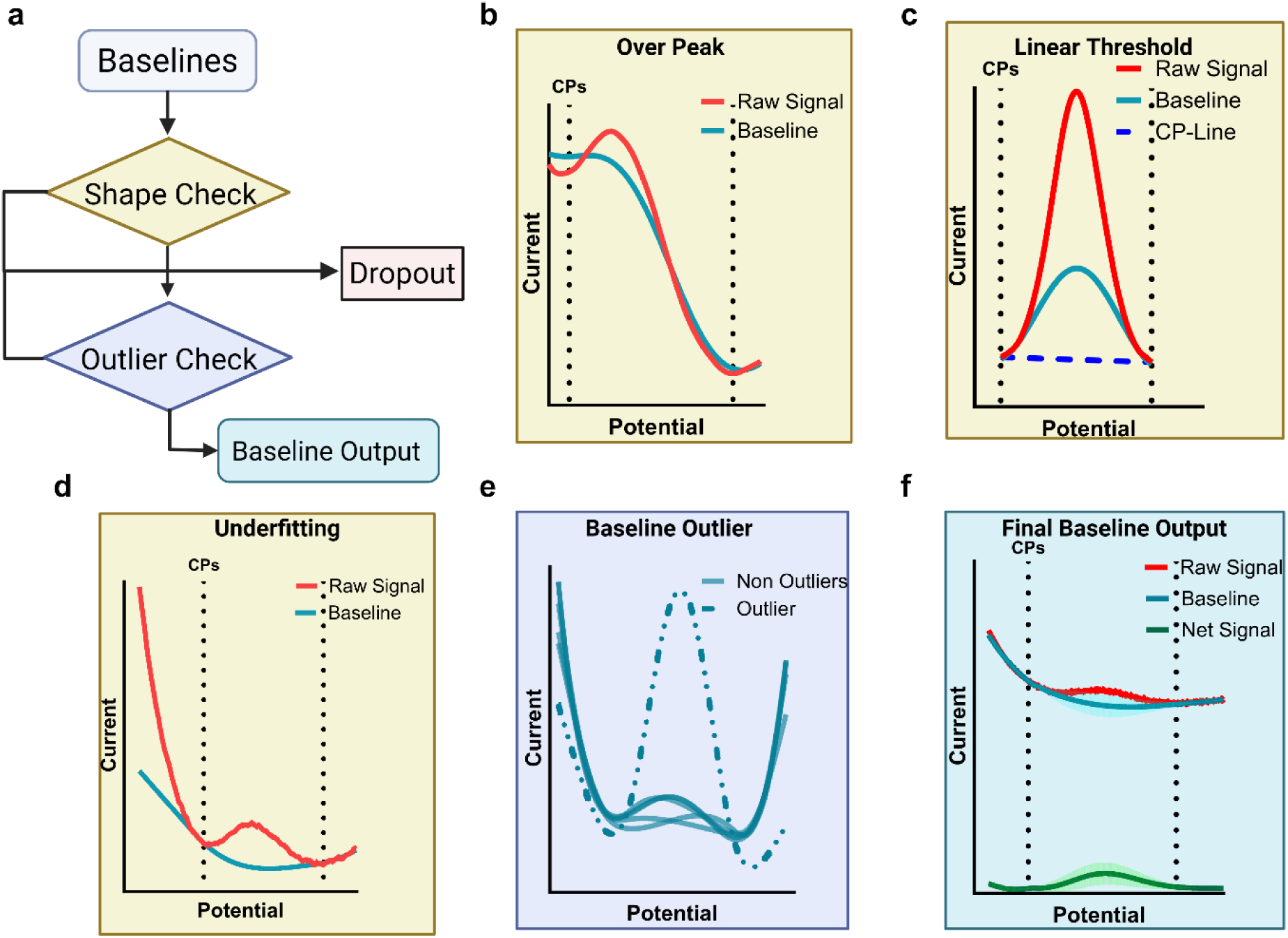
Baseline fitting analysis includes shape check and outlier check that lead to the final baseline output. **a** The screening procedure involves dropping baselines obtained from various algorithms based on shape check and outlier check. The final baseline output is resulted from the median of the remaining baselines with a 99% confidence interval. **b** Example of baseline dropout in which has more than 10% of the data points are above the raw data in the peak region. **c** Example of baseline dropout with unexpected peaks in the peak region and an AUC that differs by more than 30 % from that of a linear baseline. **d** Example of baseline dropout with low consistency (MSWE larger than 0.1) compared to the raw data in the baseline region. **e** Example of one baseline outlier using the local outlier factor algorithm. **f**. Example of final baseline output, including the median values and the 99% confidence interval from the passed baselines.

A-PACE then would bypass the AUC comparison and proceeded directly to the next step. Third, the fitted baseline was compared with the raw baseline through normalized weighted mean square error (NWMSE) defined in the **Methods**. The fitting algorithms that resulted in high errors (NWMSE > 0.1) were removed from further analysis (**Figure 3d**). After shape check, we then mitigate the negative impact of certain fitted baselines that meet the above three criteria yet exhibit extreme characteristics by applying the local outlier factor algorithm to all qualified baseline fittings and excluding outliers (**Figure 3e**). Finally, the selected baseline fittings were determined as the median of all non-outlier baselines, accompanied by a 99% confidence interval (CI) (**Figure 3f**). Compared with average value, the median value helps prevent the superposition of multiple baseline statistical instability, resulting in smoother time-series processing outcomes. And the 99% CI provides a reasonable range for peak height. We then identified a generalized and optimal A-PACE algorithm set—comprising both the CPD algorithms (search method and cost function) and baseline fitting algorithms, as shown in **Figure 4a**. First, we selected 30 proper baseline fitting algorithms (the full list in **Table S2**) from *pybaselines* package^28^ and then tested them with all 2,000 SWV curves. The peak height for each SWV curve was extracted using the screening and filtering process mentioned in **Figure 3a**. Two key metrics, success rate (SR) and normalized mean square error (NMSE) were used to quantitatively assess and compare the performance of each algorithm (more details in the **Supplementary Note 3**). To balance the SR and NMSE, we selected the top 5 baseline fitting algorithms with the highest SRs and the top 5 with the lowest NMSEs. As a result, the selected 10 baseline fitting algorithms out of 30 are shown in **Table S3**. Thereafter, the selected 10 baseline fitting algorithms can result in 1023 combinations (2^10^-1), which together with 2 CPD algorithms (BottomUp-rank and Dynp-rank as the top 2 from CPD section) can lead to a 2046 algorithm set for peak height analysis and the following screening performance evaluation. The SR and NMSE of each algorithm set are shown in **Figure 4b**. Due to complementarity among multiple baseline fitting algorithms, most algorithm sets have a SR of 100% when using the top two CPD algorithms for CPs. Since slight variations in CP positions led to different top baseline fitting algorithms for combinations (**Table S3**), baseline fitting algorithm sets with different CPD algorithms have clearly distinguishable outcomes in terms of final SR and NMSE metrics. To achieve a balance between SR and NWMSE, we calculate their z-scores for further analysis and then adopted the Markowitz efficient frontier theory to define a user utility function that allows the user to choose the best pairs of SR and NMSE from the Markowitz curve (more details in **Methods**).^29^ For example, based on a series of weightings assigned to SR, users can select weight for SR ranging from 0 to 1 with a step of 0.01 and obtain the optimal algorithm set with proper SR and low NMSE. Within the 101 possible weights, 92 weights share the same optimal algorithm set. Therefore, we listed it as the default A-PACE algorithm set. This default algorithm (marked in **Figure 4b** with a detailed algorithm list in **Table S4**) has a SR of 100% and a NMSE of 0.0196 for 2,000 SWV curves, which shows its capacity to reliably quantify peak heights across a diverse dataset and to generate independently derived time-series trajectories that exhibit requisite smoothness and coherence. SR and NMSE values from 40 folders, determined with the default A-PACE algorithm set, are shown in **Figure S5**. One thing to emphasize, even for the EAB sensors with tens of micron footprint and single-digit nanoampere current, this algorithm set has a SR of 100% and a NMSE of 0.0245 on average, which further demonstrates that A-PACE lowers the barrier to implement miniaturized biosensors for continuous monitoring.

**Figure 4.**
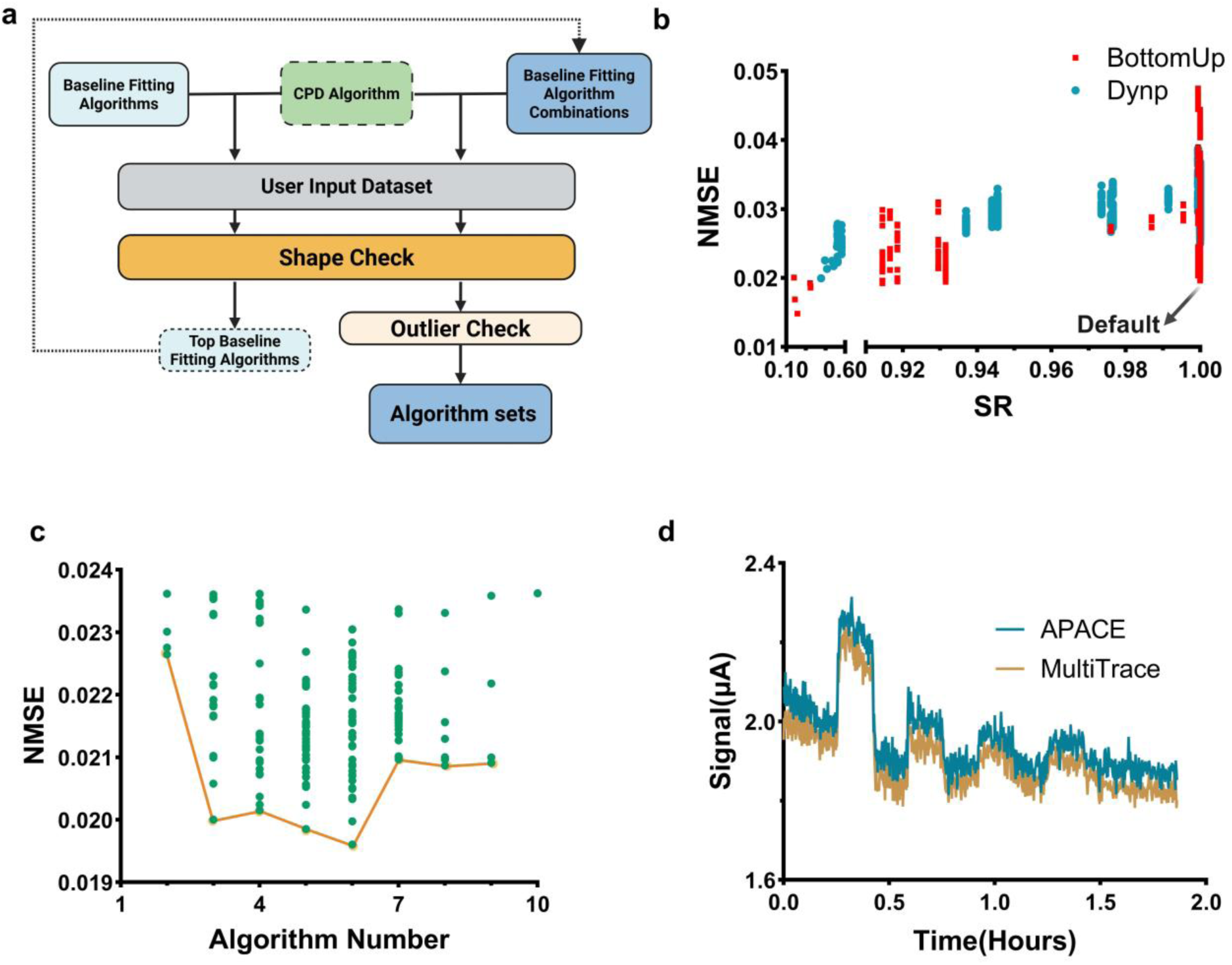
Selection process of baseline fitting algorithm sets and its performance are evaluated by success rate (SR) and normalized mean square error (NMSE) **a** Workflow of baseline fitting algorithm set selection. Users can select the corresponding algorithm set from the algorithm sets based on the input success weight, and use the time series dataset to obtain an optimized algorithm set following the same procedure, respectively. **b** SR and NMSE of 2046 algorithm sets are shown. Although two CPD algorithms produce different fitting outcomes, 1360 out of 2046 algorithm sets achieve 100% SR. The default algorithm set is obtained based on the set with one of the highest SR and one of the lowest NMSE, according to Markowitz efficient frontier theory. **c** NMSE values for the top 200 algorithm sets initially decrease with an increase in the algorithm number, followed by a subsequent rise over the algorithm number of 6. **d** Time series data obtained using the A-PACE default algorithm set is compared with the data obtained using MultiTrace. A cosine similarity of 0.9999 indicates a high level of consistency between the two tools for *in vitro* continuous electrochemical analysis.

Furthermore, we explored how the number of baseline fitting algorithms in each A-PACE algorithm set affect the SR and NMSE. We analyzed the top 200 algorithm sets with 100% SR and ranked them by lowest NMSE value (**Figure 4c**). Initially, increasing the number of algorithms improved stability (a sharp decline of NMSE from two algorithms to three algorithms) due to multiple choices of baseline fitting algorithm to cover a broad range of SWV curves in the dataset. However, once the number of algorithms exceeds a threshold (six for this dataset), interference and competition among the baseline-fitting algorithms will induce the instability of overall fitting performance (More details in **Table S5**). Therefore, the optimal number of baseline fitting algorithms was determined to be six for the default A-PACE algorithm set.

As proof of concept, this default algorithm set was applied for continuous electrochemistry in the flow cell experiment, compared to one of commercially available electrochemical software, such as MultiTrace from PalmSens, and shown almost identical real-time response with a cosine similarity of 0.9999 (**Figure 4d** and more details in **Methods**). We also validated the performance of the default algorithm set on SWV curves with different signal levels and shapes, like large, small, noisy, tilted curves as summarized in **Figure S4**. By applying different levels of Savitzky-Golay (SG) filter, all curves are reasonably smoothed and fitted. As the control group, the large SWV curve has a bell shape, similar to a Gaussian distribution (**Figure S4a**), whose fitted baseline closely matches the baseline region of the raw data with a 99% CI covering the baseline fittings from different algorithms. The noisy SWV curve (may be caused by small sensor size) as shown in **Figure S4b** is close to the bell-shaped curve in **Figure S4a** with a clear fitting after smoothing. For the small curve (**Figure S4c**) and the tilted curve (**Figure S4d**) which may be caused by biofouling, unreasonable baselines have been filtered by shape check that leads to a selected baseline with reasonable fittings. More importantly, when the default A-PACE algorithm set is suboptimal for new electrochemical data, the user can follow the same optimization pipeline (**Figure 3a**) for CPD and baseline algorithm screening to obtain a specific A-PACE algorithm set tailored to the dataset.

### Data interaction and visualization workflow

We designed a user-friendly GUI for A-PACE that streamlines data processing and visualization. The GUI workflow is illustrated in **Figure 5a**. After launching, the GUI appears in a web-based interface with two modes, real-time analysis mode and post-data processing mode, both of which are available. The real-time analysis mode enables real-time monitoring and interpretation of time-series data when measuring. This allows users to track electrochemical peak height over time during the measurement. To address instability due to fluctuating CPs over a real-time measurement, a sliding window with self-adjusting width was introduced to stabilize the real-time output (**Figure 5b**). Since the SWV peak positions are expected to be less prone to change than peak heights in a time-series data, this sliding window approach can better mitigate outlier data point issue observed in some real-time measurements. And the real-time output is continuously updated with adjusted peak heights and associated confident intervals (**Figure 5c**). The post-data processing mode is designed to maximize computational efficiency in analyzing large datasets collected across various time periods, concentration ranges, and electrochemical parameters, e.g., amplitude, frequency, potential step and channel (more details in **Figure S6**). This mode enables the user to efficiently organize, edit, and analyze the dependency of experimental variables on the measured signals. **Figure 5d** shows the summary table of measurement information and **Figure 5e** highlights the 3D plotting that visualizes the relation of different variables.

**Figure 5.**
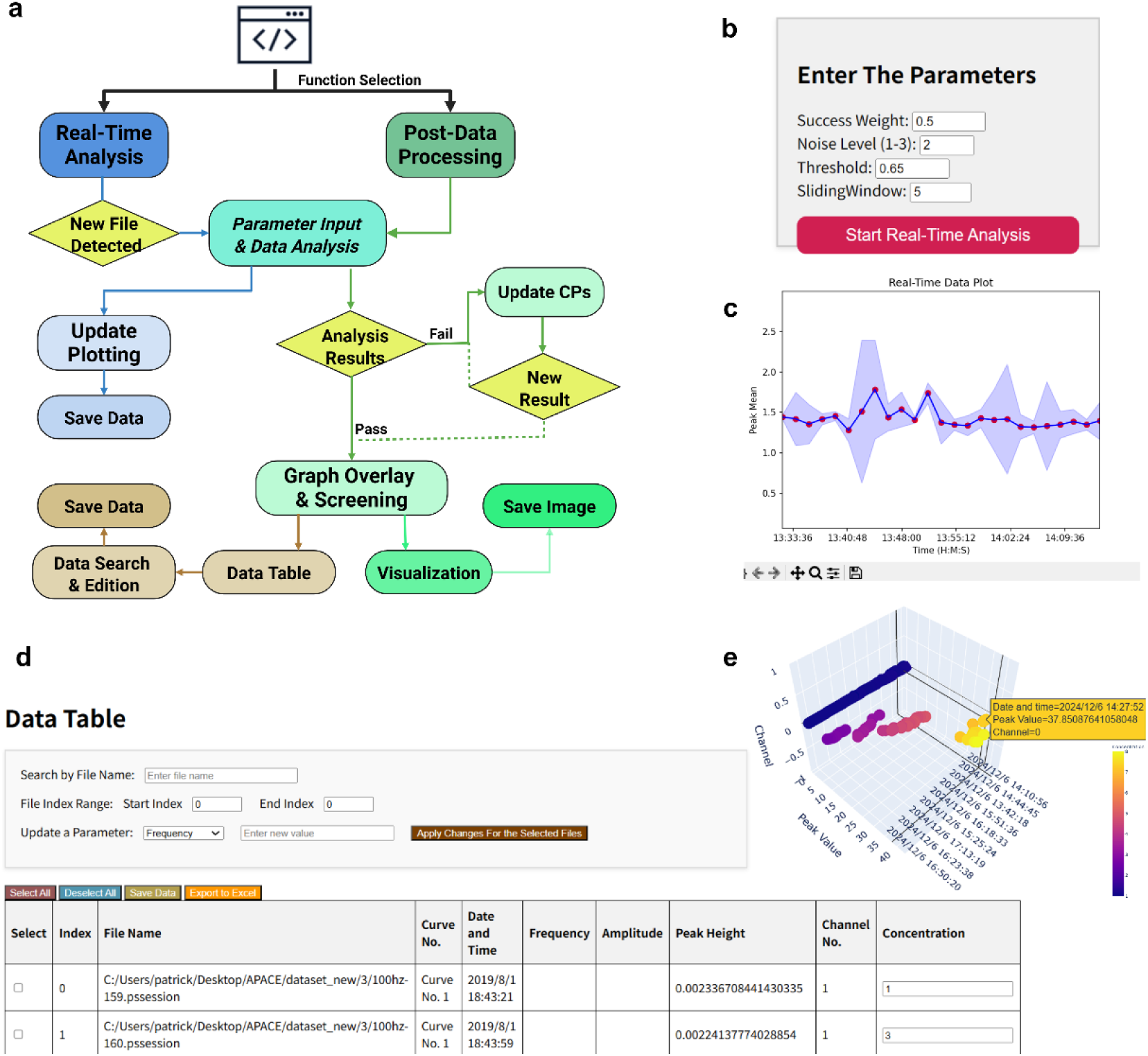
GUI workflow and functions: Real-time analysis mode offers fast monitoring of continuous electrochemical signals, and post-data processing mode provides fine tune parameters, customized data output format, interactive data visualization. **a** GUI workflow that shares similar algorithm pipelines for real-time analysis mode and post-data processing mode, enabling users to adjust the analysis steps with various options for spreadsheet and figure output. **b** Parameter input interface that allows the user to adjust success weight, noise level, peak region threshold and sliding windows size. **c** An example of the real-time analysis window showing the time series results. The sliding window offers an option for real-time analysis mode to obtain more stable signal analysis. **d** Data table contains electrochemical metadata information for user review, editing, and output, including the ‘Curve path and index’, ‘Data and Time’, ‘Frequency (Hz)’, ‘Amplitude (V)’, ‘E Step (V)’, ‘Peak Height (*µ*A)’, ‘Channel Number’, and ‘Concentration’. **e** An example of a 3D graph based on ‘Data and Time’, ‘Peak Value’, and ‘Channel Number’ information. Various options for figures can be generated based on the properties selected from the data table shown in panel **d**.

### Case studies

Typically, EAB sensors exhibit limited signal longevity—remaining reliable for less than one day *in vivo* and seldom lasting beyond one week *in vitro*^30–32^. This short signal lifetime significantly makes systematic studies time-consuming and labor-intensive, creating a major bottleneck to the advancement of EAB technology. As demonstrated earlier, A-PACE enables high-throughput, objective analysis with consistent accuracy. In the following case studies, we highlight A-PACE’s performance using two challenging test datasets: (1) Month-long electrochemical data in serum and (2) Week-long electrochemical data after multi-day intravenous implantation in free-moving rats.

#### Month-long electrochemical data in serum

To assess A-PACE’s ability to process SWV curves under severe signal degradation, we conducted a month-long *in vitro* measurement using a kanamycin-sensitivity EAB sensor with no surface protection, incubated in undiluted human serum (**Figure 6a**). To evaluate its performance across a broad analyte concentration range, electrochemical data were collected on days 0, 1, 3, 7, 14, and 28, with kanamycin concentrations ranging from 10 µM to 10 mM. A total of 288 SWV curves were processed in just 18.7 seconds. The representative results are shown in **Figure 6b-g**. From day 0 to day 14, the SWV signal baselines became increasingly tilted, and the peak responses progressively less distinguishable. This signal decay over days is inevitable in biosensing monitoring in serum mainly due to non-specific protein adsorption and the loss of aptamers attached to the electrode. Correspondingly, wider confidence intervals are observed, although A-PACE can consistently estimate the signal baselines from raw SWV curves. By day 28, signals were highly distorted and dominated by noise, rendering manual annotation of baselines and peak heights nearly impossible. Remarkably, A-PACE continued to extract CPs and export baselines for peak height analysis, which can pass the restrict screening and filtering process (more details in the **Supplementary Note 4**). As shown in **Figure 6h**, A-PACE effectively tracks signal degradation over a four-week period, successfully extracting peak height as low as 13.6% of the original peak height from these continuously decreased electrochemical signals. These results demonstrate A-PACE’s robust performance and highlight its capability to handle small and noisy electrochemical signals from long-term *in vitro* experiments.

**Figure 6.**
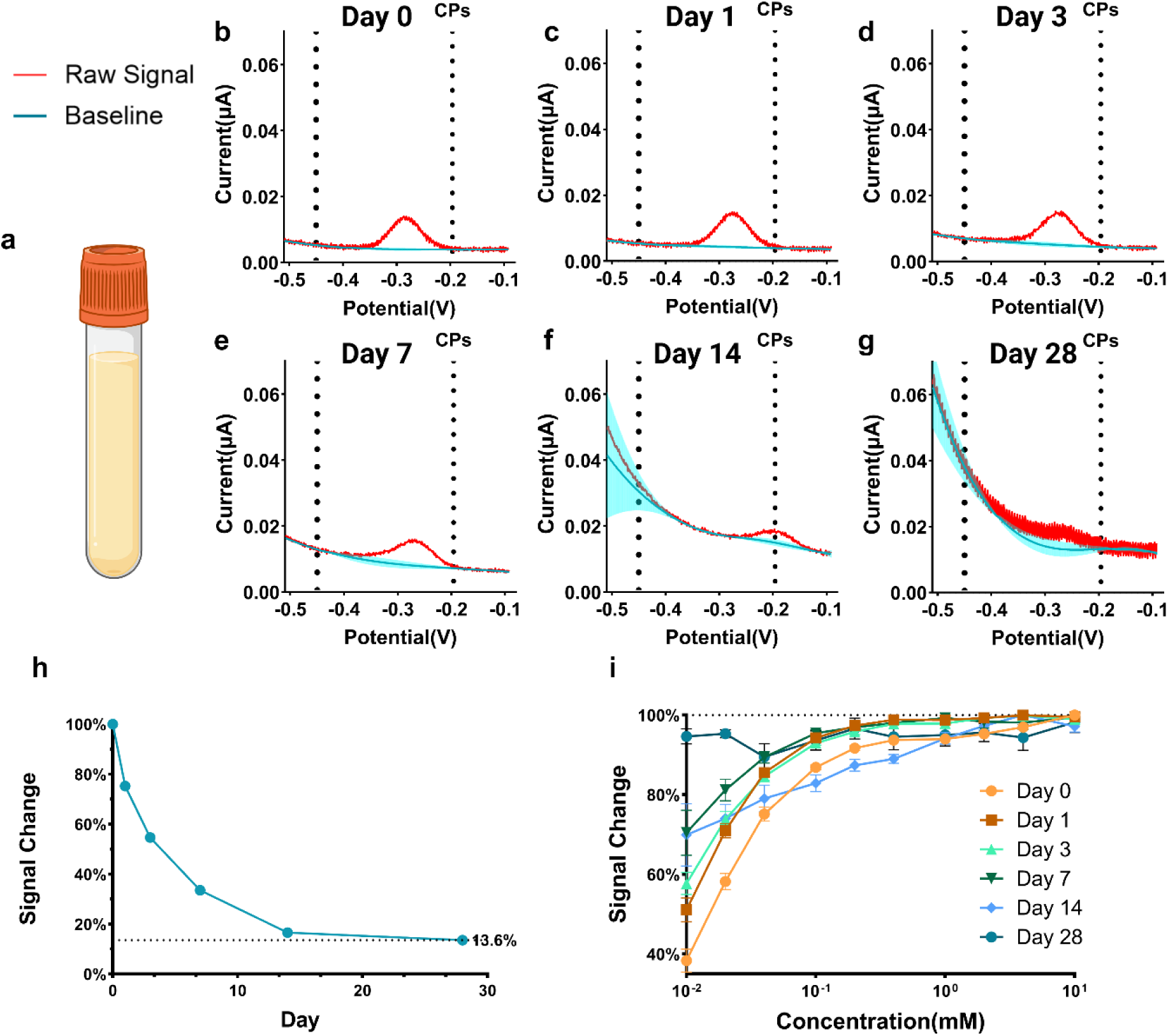
Month-long analysis of human serum samples. Over the 28-day period, A-PACE with the default algorithm set shows stable concentration-signal responses at different days with a broad concentration range. **a** *in vitro* detection in human serum. SWV measurement was carried out over the potential range of ∼-0.5 V to ∼0.0 V, amplitude of 20 mV, step size of 1 mV, and pulse frequency of 50 Hz. **b-g** *in vitro* signal details over 28 days. Even though the signals became increasingly tilted with decreasing SNR over 28 days, A-PACE can find baselines for all cases with peak height extracts for concentration-signal response analysis. **h** *in vitro* signal change over 28 days with the final signal intensity declined to 13.6% of its initial value. **i** *in vitro* concentration-signal responses over 28 days. Even on Day 28, the overall signal trend from A-PACE analysis remained consistent, demonstrating its capacity for month-long *in vitro* analysis.

Furthermore, to evaluate A-PACE’s capability in analyzing signal variations induced by concentration changes, we applied it to generate calibration curves for long-term kanamycin monitoring. Kanamycin was incrementally spiked up to total concentration of 10 mM, and signals were recorded at each concentration level on days 0, 1, 3, 7, 14, and 28 using four channels (more SWV fitting results are summarized from **Figure S7** to **Figure S12**). From day 0 to day 7, normalized signal intensities increased consistently with rising kanamycin concentrations. By day 14, the upward trend persisted, although an anomalously high response was observed at 0.04 mM. By day 28, the calibration curve exhibited substantial fluctuations, which were attributed primarily to measurement variability rather than limitations in A-PACE’s signal analysis. Importantly, A-PACE achieved a 100% SR in processing all signals across the 28-day kanamycin sensitivity analysis. The analysis showed no extreme outliers, and a general trend of increasing signal with concentration was still discernible even after the device incubation in undiluted human serum for 28 days. For comparison, **Figure S13** presents signal degradation analysis using the auto-detect peak function from MultiTrace software. Although MultiTrace exhibited a similar degradation trend with time((**Figure S13a**) and concentration((**Figure S13b**,), it failed to analyze 38 signals out of 40 on day 28 (**Figure S13b**, signals with zero value represent failed cases), resulting in an apparent signal intensity of only 2.20% of the original value (**Figure S13a**). Furthermore, while both A-PACE and MultiTrace performed comparably in detecting concentration-dependent signal increases during the first 14 days, MultiTrace exhibited high variability across different channels (electrode replicates), leading to large error bars and inconsistent results. This indicates that data processed with MultiTrace exhibits substantial variability arising from the data processing tool itself, leading to significant unreliability in the final results. In contrast, A-PACE maintained small error bars throughout the entire 28-day period (**Figure 6i**), indicating high precision and reliable performance across all channels under conditions of signal degradation and electrode variability.

#### Week-long electrochemical data in rats

Since the default algorithm set was trained using the electrochemical data from *in vitro* measurements, we evaluated its performance using ten short period *in vivo* datasets, practicing the scenario for real-time molecular monitoring with no prior information about individual patients and sensors. For comparison, the results were also analyzed using A-PACE and MultiTrace. As shown in **Figure 7a**, both methods captured signal changes following target addition and A-PACE shows less variation of peak height estimation from all-time series data. In **Figure 7b**, MultiTrace failed to process three SWV curves and produced two outlier results, whereas A-PACE maintained a 100% SR and delivered stable, consistent signal outputs. Eight additional examples are provided in **Figure S14**. And cosine similarities are also calculated across all ten cases in **Table S6**. Seven out of ten cases have a cosine similarity over 0.95, which shows that both tools have similar general trend of analysis results. However, across all ten cases A-PACE maintained all 100% SR with relatively smoothed results than MultiTrace, and clearly resolved signal changes in most target addition events—capabilities not achievable with existing electrochemical analysis tools. To further explore the differences in SR between the two tools, we present in **Figure S15** the analytical results obtained by both tools at the failed points of MultiTrace from **Figure S14g**, along with their preceding and subsequent time points. Two tools share the same CPs before and after the failure time point, which led to a similar data trend. However, when MultiTrace failed to identify the CPs, A-PACE can still successfully locate the CPs and proceed with the following analysis. Moreover, the CPs identified by A-PACE closely matched those at adjacent time points, resulting in smoother and more consistent time-series analytical outcomes.

**Figure 7.**
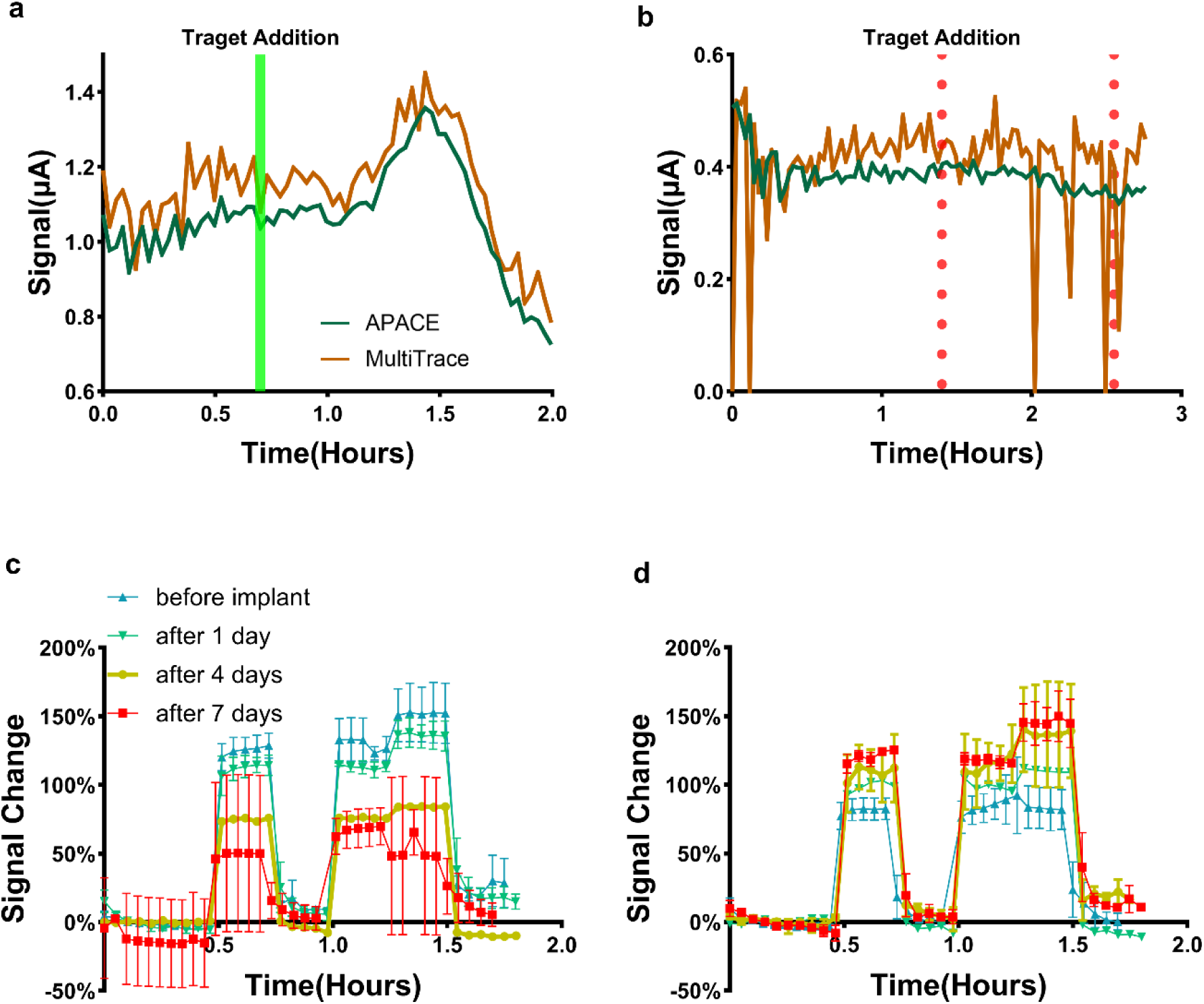
*in vivo* analysis performance of A-PACE. The A-PACE with the default algorithm set shows better performance than the MultiTrace in *in vivo* time series data. In the analysis of *in vivo* intravenous blood, the default algorithm set based on the *in vitro* dataset has relatively large signal variation on day 7. In contrast, the optimal algorithm set based on the *in vivo* data under analysis provides consistent time and concentration response throughout week-long period. **a** Data processing results of the *in vivo* signal with clearly added target signals. Both tools detect the target addition point (green solid line) with A-PACE has smoother results. **b** Data processing results of the *in vivo* signal with ambiguous target addition points (red dot line). Neither tools can find them, and MultiTrace has more failed cases. **c-d** Real time kanamycin monitoring after multi-day intravenous implant. **c** A-PACE with the default algorithm set that analysis results have a larger error bar with 6.2% failed cases only on day 7. **d** A-PACE with the optimized algorithm set that analysis results have all 100% success rates with instant drug response over 7 days.

To further evaluate A-PACE’s applicability in more complex biological environments, we conducted *in vivo* electrochemical measurements after multi-day intravenous implantation in freely moving rats using our nanostructured sensor platform^33^. The nanostructured sensors were used to continuously monitor sequential kanamycin concentrations of 1 mM, 0 mM, 1 mM, 2 mM and 0 mM in PBS buffer after multiple days of intravenous implantation in freely moving rats. These measurements were performed with different sensors on days 1, 4, and 7 following femoral vein implantation (More experimental details in **Methods**)^33^. To better illustrate the signal’s concentration-dependent response using repeated measurements at a single concentration, we computed the daily mean of the first ten signal points at 0 mM and used this value to normalize all signals. Initially, we applied unsupervised default MultiTrace built-in function to analyze these datasets, with results summarized in **Table 1**. Over the seven-day period, MultiTrace consistently demonstrated SR values lower than 70%. By day 7, it failed to process any of the SWV curves. As shown in **Figure S16**, *in vivo* signal degradation severely impacted MultiTrace performance: signal variability before implantation and on day 1 led to large error bars, compromising the reliability of results. On day 4 and day 7, the built-in mode from MultiTrace yielded low SR values and unreliable outputs, rendering the analysis uninterpretable.

**Table 1.** Success rates of MultiTrace, A-PACE (Default algorithm set) and A-PACE (Optimized algorithm set) in analyzing *in vivo* measurement results over 7 days.

| Days | MultiTrace | A-PACE-Default | A-PACE-Optimized |
| --- | --- | --- | --- |
| Before Implant | 69.5% | 100% | 100% |
| After 1 day | 54.3% | 100% | 100% |
| After 4 days | 63.9% | 100% | 100% |
| After 7 days | 0% | 93.8% | 100% |

We then applied A-PACE using real-time analysis mode with the default algorithm set (65% peak region threshold, smoothing level 2), which processed 493 SWV curves from seven-day *in vivo* SWV measurement. Compared to MultiTrace, A-PACE achieved higher SR values (more SWV fitting results are summarized from **Figure S17** to **Figure S20**). By day 4, A-PACE maintained a SR of 100%, and the small error bars in **Figure 7c** confirm the reliability and consistency of its results. By day 7, the signal distinction during transitions from 1 mM to 2 mM and subsequently back to 0 mM was no longer evident. An example of a failed case is shown in **Figure S21a**. In that instance, although the smoothed signal was distinguishable and the detected CPs were correctly located, the baseline fitting from the default algorithm failed to meet screening and filtering criteria. This failure can be attributed to the default algorithm set being trained on *in vitro* datasets, which may not be fully optimized for *in vivo* signal characteristics.

Using the same *in vivo* dataset here, we next evaluated further data analysis enhancement in the post-data processing mode, with same threshold and smoothing level. Then we applied our algorithm selection pipeline to screen for the most suitable algorithm set tailored to this *in vivo* dataset. With existing aimed data for algorithm selection, an optimized algorithm set has a better performance than default algorithm set in real-time mode. The details of post-data processing mode with the optimized algorithm set are presented in **Table S7**. This newly selected algorithm set employs different baseline fitting algorithms and successfully resolves cases that the default algorithm set could not (**Figure S21b**). With this optimized algorithm set, A-PACE achieved a SR of 100% across all seven days and produced smaller error bars, particularly on day 7 (**Figure 7d**). Moreover, the concentration-dependent signal responses are more consistent among different days, which also indicates the nanostructured biosensors are well-operated with target sensitivity unchanged after one-week intravenous implant. These improvements demonstrate that the optimized algorithm set used in the post-data processing mode not only enhances analytical performance but also extends the lifespan of implanted electrochemical biosensor other than applying conventional surface protection strategies.

## DISCUSSION

Recent research on electrochemical biosensors has concentrated mainly on material innovation and novel measurement protocols for continuous monitoring. Signal processing advances have likewise tended to produce bespoke, high-performance algorithms optimized for specific signal waveforms or devices. Developing such tailored routines can be labor-intense, demands deep domain expertise, and often yields solutions that translate poorly across applications. Unsupervised learning models promise broader applicability, yet their training generally requires extensive datasets, significant computational resources and meticulous hyper-parameter tuning. To navigate this diversity in signal categories and morphologies, we introduced A-PACE—a constrained yet adaptive, high-throughput toolkit for continuous electrochemistry. By combining change-point detection with multi-algorithm baseline fitting, A-PACE handles a wide spectrum of electrochemical outputs, including low-SNR traces that accompany aggressive sensor miniaturization. Its capacity to resolve tilted and/or distorted peaks enables clearer discernment of biofouling effects in the resulting dataset than is possible with existing analysis tools.

Beyond post-hoc analysis, A-PACE supports real-time electrochemical signal analysis, delivering immediate feedback that streamlines experimental workflows. Because the pipeline is algorithm-driven rather than operator-dependent, it mitigates subjective bias and enhances inter-lab reproducibility. Widespread adoption would thus reduce variability arising from disparate signal processing protocols, thereby facilitating comparisons across different studies. The framework of A-PACE is not without constraints. First, accurately identifying peak positions in complex, multi-peak voltammograms remains challenging; second, performance hinges on the breadth and quality of the CPD and baseline fitting libraries. These limitations, however, stem less from intrinsic shortcomings than from the sheer heterogeneity of measurement conditions. Crucially, A-PACE’s adaptive pipeline can retune itself: when presented with an unfamiliar dataset it iteratively searches its algorithm bank, selects the optimal combination, and typically achieves a high success rate with excellent stability. Its web-based GUI further allows users to tweak change point locations interactively, connecting automation with expert oversight. The entire platform is open-source and hosted on GitHub, inviting community contribution and rapid evolution.

In summary, A-PACE processed month-long *in vitro* serum data and week-long *in vivo* rat data, extending the effective analytical lifetime of electrochemical biosensors well beyond the conventional window. Additionally, this CPD and multi-baseline fitting framework is readily transferable to other biosignals, such as electroencephalogram (EEG) and electrocardiogram (ECG), where reliable peak detection and rapid signal classification are paramount. Looking ahead, the scarcity of large, high-quality electrochemical datasets limits deeper forays into machine-learning pipelines. Deploying arrayed or multiplexed electrochemical sensors that generate data at scale would unlock the full potential of A-PACE by feeding it richer training electrochemical datasets. In such scenarios, this toolkit’s rapid, standardized preprocessing becomes a critical point for automated, long-term monitoring systems. Coupled with recent advances in self-supervised learning, A-PACE could underpin closed-loop platforms in which sensors stream raw voltammograms, the software cleans and quantifies them in real time, and downstream models deliver actionable insights with minimal human intervention. A-PACE bridges the gap between specialized algorithm development and universal applicability. It offers a robust, open, and extensible solution that not only improves today’s electrochemical biosensing experiments, but also lays the groundwork for tomorrow’s autonomous, data-driven diagnostics.

## METHODS

### Materials

All chemicals were purchased from Sigma-Aldrich unless otherwise specified. Thiolated polyethylene glycol (PEG-SH, 8-arm PEG-SH, MW 20 kDa) was obtained from Creative PEGWorks. Kanamycin monosulfate, USP Grade, was sourced from Gold Biotechnology. PFA-coated tungsten wires (bare diameter 0.002”) was purchased from A-M Systems. 10k gold wire (diameter 0.01”) were purchased from Amazon. Miniature heat-shrink tubing (pre-shrink inner diameters of 0.006” or 0.01”) was obtained from Nordson Medical and McMaster-Carr. The kanamycin aptamer was synthesized by Integrated DNA Technologies, Inc. (IDT) with the following sequence: /5ThioMC6-D/GGGACTTGGTTTAGGTAATGAGTCCC/3MeBlN/, where 5ThioMC6-D denotes a thiol modification at the 5’ end and 3MeBlN indicates a methylene blue (MB) label at the 3’ end. Phosphate-buffered saline (PBS) used in this study was prepared by diluting 10X PBS and MgCl_2_ stock solutions with nuclease-free water (NFW) to a final concentration of 1X PBS containing 2 mM Mg^2+^.

### EAB sensor preparation

#### Planar gold and nanoporous gold fabrication

A schematic of the device fabrication process is as described in our prior work^33^. For *in vitro* sensors, a 90 nm-thick gold (Au) layer was deposited onto pre-cleaned glass slides using electron beam evaporation (ATC-E, AJA International) through a shadow mask to define 1 mm × 1 mm working electrodes. A 10 nm titanium (Ti) adhesion layer was applied prior to Au deposition. For *in vivo* unprotected microwire sensors, gold wires were cut into 6 cm lengths and used without further processing. For *in vivo* nanostructured sensors, nanoporous gold microwires were prepared by etching 10k yellow gold wire in 70% (v/v) nitric acid for 15 minutes, which is further treated as described in our prior work^33^. The microwire wires were insulated with heat-shrink tubes around the body of the wires, with a ∼2–5-mm length of electrode exposed for aptamer functionalization, as described in our prior work^34^. All electrodes were sequentially sonicated in acetone, 70% isopropyl alcohol, and nuclease-free water (NFW) for 2 minutes each, then stored in NFW prior to functionalization.

#### Sensor functionalization

Aptamer switches were suspended in NFW at a final concentration of 100 µM and stored at -20 °C before usage. To produce free thiol groups for aptamer immobilization, a 1,000-fold molar excess of freshly prepared Tris(2-carboxyethyl) phosphine hydrochloride (TCEP) solution was added to the aptamer switch solution and reacted for 40 min at room temperature in dark. The mixture was then diluted to 1 µM aptamer in PBS. Electrodes were briefly dried with N_2_ gas. For *in vitro* experiments, a polydimethylsiloxane (PDMS) chamber was prepared by punching a 6-mm-diameter hole in the 5-mm-thick PDMS film and subsequently putting the PDMS film onto the glass slide, such that the Wes were confined within the chamber. For *in vivo* experiments, the microwire electrodes were incubated in tubes. The samples were then immersed in the 1 µM aptamer solution for 1 hour. After immobilization, electrodes were washed twice with PBS. For some sensors, polymer coating may be used by incubating the aptamer-immobilized electrode with PEG solutions at room temperature for 3 hours. PEG was prepared by dissolving in 70% isopropyl alcohol (1 wt%) with 65 °C heating or sonication. The electrodes were then washed with excess PBS twice and incubated with ∼10 mM 6-mercapto-1-hexanol (MCH) for 3 hours at room temperature to fully passivate the remaining electrode surface. The sensors were washed in excess PBS twice and then stored in PBS buffer at room temperature before electrochemical measurement.

### Electrochemical dataset acquisition

#### In vitro electrochemical measurements

Electrochemical measurements were conducted with PalmSens 4 potentiostat (PalmSens) and EmStat4 (PalmSens) with multiplexer (MUX8-R2 or MUX8, PalmSens). For *in vitro* experiments, the measurements were performed in PDMS wells. Commercial Ag/AgCl reference electrodes and Pt wire counter electrodes were used as received from CH Instruments. SWV was carried out in buffers or human serum over the potential range of ∼-0.5 V to ∼0.0 V with an amplitude of 20 mV, step size of 1 mV, and pulse frequencies ranging from 20 to 200 Hz. To measure biosensor sensitivity *in vitro*, we obtained dose-response curves by subjecting the sensors to kanamycin concentrations ranging from 0.01–10 mM in various biological matrices by spiking in kanamycin stock solution (dilution < 1% v/v) at room temperature. Long-term incubation was achieved by loading 100–200 µL undiluted biofluids into the PDMS chamber, which was temporarily sealed with a PDMS lid before next measurement. For chronic measurements, biofluids were aliquoted from the same batch of biological matrix and thawed 2 hours before the measurement followed by gently mixing.

#### Live animal electrochemical measurements

Live animal studies were performed with male Sprague–Dawley rats under Stanford Laboratory Animal Care (APLAC) protocol number 33226. All rats used in this work were purchased from Charles River Laboratories, with a weight range of 200– 450 g and an age range of 1–4 months. The rats were anesthetized using isoflurane gas (2.5%) during implantation procedures and monitored continuously. The femoral veins of the rats were isolated and exposed with a small incision, through which the microwire probes were implanted and pushed far into the vein. The microwire probes were secured in the place while maintaining the intravenous blood flow, after which the incision was closed and sutured with absorbable sutures and a small amount of tissue glue to anchor the suture wire. The microwire probes were sterilized with a UV light sanitizer (Amazon) prior to implantation. The implantation wounds were sutured and secured with wound clips. The animals were monitored daily for post-surgery recovery and maintained for up to seven days. At the end of the experiments, animals were anesthetized using isoflurane gas and the microwire probes were retracted for post-implantation analysis. Rats were then euthanized by exsanguination while under general anesthesia.

### A-PACE Platform

#### Change point detection

We evaluated different smoothing levels on signals of varying Signal-to-Noise Ratios (SNR), demonstrating that lower smoothing was sufficient for high-SNR signals (>40), while higher smoothing levels were essential for effective noise removal in low-SNR signals (<10). CPD was implemented using the Python package Ruptures^27^, testing four search methods-Dynamic programming (Dynp), Binary segmentation (Binseq), Window sliding segmentation (Window) and Bottom-up segmentation (BottomUp) combined with six different cost functions-CostL1 (l1), CostL2 (l2), CostNormal (normal), CostRank (rank), CostCosine (cosine) and CostRbf (rbf). Performance was assessed by comparing algorithm-detected change points (CPs) to manually annotated CPs, considering consistency within a 10% deviation threshold. BottomUp-rank and Dynp-rank algorithms achieved the highest accuracy (>80%), demonstrating strong alignment with human annotation. The dataset included signals measured at frequencies from 50 Hz to 400 Hz, with current levels ranging from nanoamperes (nA) to microamperes (*µ*A).

#### Baseline fitting and screening

Baselines underwent rigorous shape verification, with thresholds set to exclude those significantly deviating from the raw curve or peak region. MSE (mean square error) is ideal as an objective measure of model performance^35^. Since data points near the peak region critically determine the baseline in this area and significantly influence the final peak height, increased weighting was applied to this region during the evaluation of fitting results.

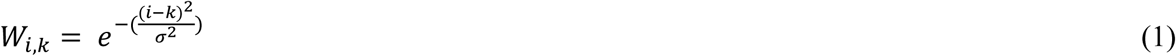

where *i* is the index of data point while *k* is the index of one CP. And *σ* =(0.25∗*L*)^2^, where *L* is the total data length of baseline region.

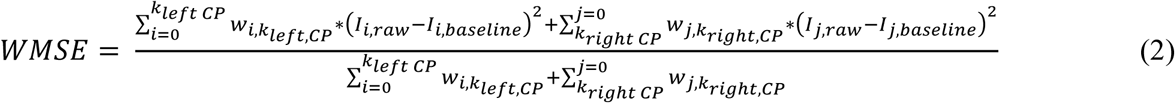

where *k_left CP_* and *k_right CP_* are the index of left and right CPs, respectively. And *I_raw_* is the raw signal while *I_baseline_* is the baseline data. To balance different signal strength levels (from 1 nA to 1 *µ*A) within the dataset, the WMSE for each curve is also normalized with the signal range:

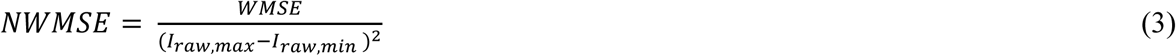

where *I_raw,max_* and *I_raw,min_* are the maximum and minimum raw signal within baseline region, respectively. Then the final NWMSE is used to define alignment between the raw signal and each fitted curve.

After screening algorithms against 2,000 experimental curves, optimal baseline algorithm sets were identified based on SR and NMSE metrics, then exhaustively combined and further refined using outlier detection techniques. To balance fitting accuracy and stability, standardized SR and NMSE (Z-scores) were computed for the determination of the most suitable algorithm combinations according to user preference:

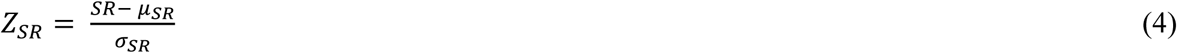

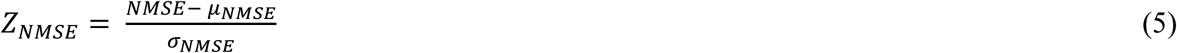

, where *µ* and *σ* are mean value and standard distribution throughout 40 folders, respectively. And a weight for *Z_SR_* was applied for algorithm set selection.

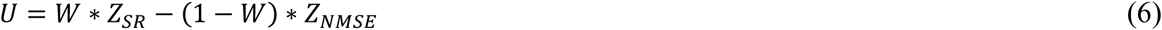

where *W* represents user’s preference to success rate.

The cosine similarity was calculated to compare the shape similarity between A-PACE’ time series results and MultiTrace’s:

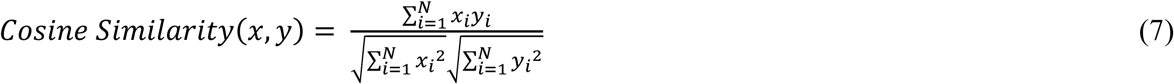

where *x_i_* and *y_i_* represent the peak heights of A-PACE and MultiTrace at time point *i*, respectively. All tests and experiments were carried out on a computational platform with an Intel Core i7-12700 CPU (12 cores, 20 threads) and 128 GB of DDR5 RAM (3600 MHz).

#### Data input, processing, and visualization

The developed GUI significantly enhances laboratory data analysis workflows by offering an accessible, intuitive platform for real-time and post-experiment data processing, including file extraction, organization, and comprehensive visualization in 2D and 3D. Leveraging Flask, Tkinter, NumPy, SciPy, matplotlib, and Plotly, it facilitates both automated and user-driven interactions, allowing researchers to dynamically adjust critical parameters such as peak height, frequency, and concentration through intuitive controls. Key features include batch processing, dynamic integration of custom algorithms, robust error handling, export functionalities, and accessibility enhancements such as keyboard shortcuts and screen reader compatibility, designed to accommodate users of varying expertise levels. For post-experiment analysis, the interface efficiently utilizes multi-core processing for CPD and baseline fitting, categorizing outcomes clearly into “Pass” or “Fail” groups, with options for user-driven refinement of analysis results. Future expansions aim to integrate cloud storage, collaborative features, and advanced machine learning capabilities, setting a new benchmark for computational tools in experimental research environments.

## Supporting information

https://github.com/ND-FULAB/A-PACE.git

## Data availability

Data supporting the findings of this study are available within Supplementary Notes, and additional data are available from the corresponding author upon reasonable request.

## Code availability

The code for A-PACE is available at https://github.com/ND-FULAB/A-PACE.git. The other codes written for and used in this study are available from the corresponding author upon reasonable request.

## Acknowledgements

This work was supported by internal funds from the University of Notre Dame. The experimental data collected and analyzed in this work were supported by the National Institutes of Health (NIH, OT2OD025342). Figures created in BioRender.

## Author contributions

Y.J. and K.X.F. conceived the idea of A-PACE and designed the project workflow. K.X.F. supervised the project. Y.C. and K.X.F. conducted the experiments and collected the electrochemical data. Y.J. implemented the core functions of A-PACE and analyzed the data. Y.Cai and J.L. advised on data analysis. K.Z. designed the GUI. H.J.O. contributed to an early version of the electrochemical data analyzer, which was later replaced by A-PACE. Y.J., K.X.F., and Y.C. co-wrote the paper. All authors discussed the results and commented on the paper.

## Additional information

