## Supplementary material for "Algorithm-Powered Analyzer for Continuous Electrochemistry: A Toolkit for Real-Time Electrochemical Data Analysis": https://github.com/ND-FULAB/A-PACE.git

### 1 Supplementary Note 1—Change Point Detection

#### 1.1 Signal Smoothing Process and Effect of Smoothing Level

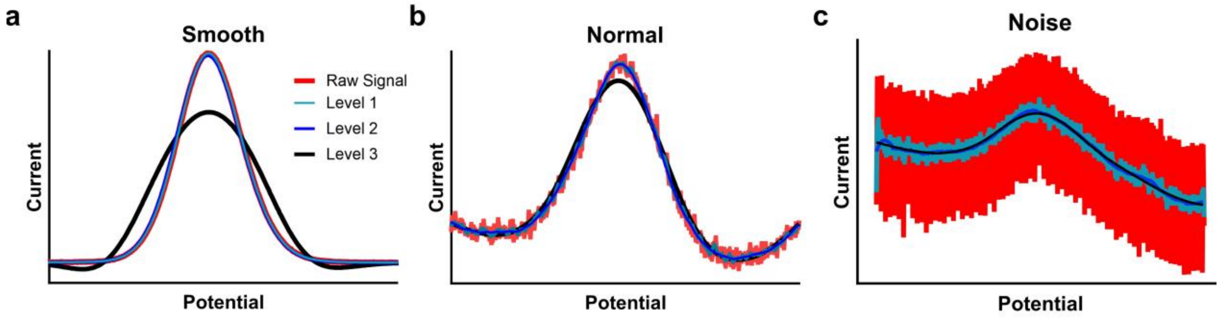

**Figure S1.** Example of curves with different Signal-to-Noise Ratio (SNR). **a** An example of curves with high SNR (over 40). Level 1 smoothing obtains perfect smoothing result. Level 2 smoothing results in a slight decrease in peak height, while Level 3 completely alters both the peak height and shape. **b** An example of a curve with normal SNR (between 15 and 40). Level 2 smoothing obtains a clean result. Level 1 smoothing remains noise, while Level 3 alters both the peak height and shape. **c** An example of curves with low SNR (under 15). Only level 3 smoothing effectively removes noise while preserving a smooth, usable signal. Multiple layers of Savitzky-Golay (SG) filters are applied for signal smoothing. Level 1 (for SNR < 15) includes a single layer with a SG smoothing window of 2%. Level 2 (for 15 < SNR < 40) applies SG filter with a double-layer smoothing windows of 2% and 10%. Level 3(for SNR > 40) uses triple layers with windows of 2%, 10% and 33%. And the default smoothing level is set as 2.

#### 1.2 Introduction to *Ruptures*

The CPD was carried out with Python Package Ruptures[1]. *Ruptures* is a Python library for change point detection. This package provides methods for the analysis and segmentation of non-stationary signals. Implemented algorithms include exact and approximate detection for various parametric and non-parametric models.

**Search Methods**: In this study, we applied and tested Dynamic programming (Dynp), Binary segmentation (Binseg), Bottom-up segmentation (BottomUp) and Window sliding segmentation (Window).

**Cost Functions**: For each search method mention above, we applied six cost functions: least absolute deviation (l1)[2], least squared deviation (l2), Gaussian process change (normal)[3,4] , Rank based change (rank)[5] and two kernelized functions with cosine similarity(cosine)[6–8] and radial basis function kernel (rbf)[8,9].

**Constraint**: To find the most proper change points for peak and baseline region segment, we preset the number of change points as three. The middle one is peak vertex while the rests are for following baseline fitting. Meanwhile, the subsample size was set as 1 for accuracy.

#### 1.3 Dataset Details

**Figure S2a** shows the details of CPs from BottomUp-rank. Compared with human labeled CPs, the detected CPs show good consistency, which indicates that such CPD method aligns with human subjectivity. **Figure S2b** and **Figure S2c** further show the details of dataset. Among 40 folders, the measurement frequency includes 50 Hz, 100 Hz, 200 Hz, and 400 Hz. And the current level of signals ranges from nA to $\mu$A. Detailed information of folders is shown in **Table S1**.

| Folder Index | Current Level | Frequency (HZ) | Amplitude (V) | E Step (V) | Labeled CP left (V) | Labeled CP right (V) |
|---|---|---|---|---|---|---|
| 1 | nA | 200 | 0.05 | 0.001 | -0.40 | -0.20 |
| 2 | nA | 200 | 0.05 | 0.001 | -0.40 | -0.20 |
| 3 | nA | 100 | 0.05 | 0.001 | -0.42 | -0.20 |
| 4 | nA | 100 | 0.05 | 0.001 | -0.39 | -0.23 |
| 5 | nA | 200 | 0.05 | 0.001 | -0.39 | -0.20 |
| 6 | nA | 200 | 0.05 | 0.001 | -0.39 | -0.20 |
| 7 | nA | 200 | 0.05 | 0.001 | -0.39 | -0.20 |
| 8 | nA | 200 | 0.05 | 0.001 | -0.40 | -0.19 |
| 9 | µA | 100 | 0.05 | 0.001 | -0.43 | -0.20 |
| 10 | µA | 200 | 0.05 | 0.001 | -0.40 | -0.29 |
| 11 | µA | 200 | 0.05 | 0.001 | -0.40 | -0.19 |
| 12 | µA | 100 | 0.05 | 0.001 | -0.40 | -0.20 |
| 13 | µA | 200 | 0.05 | 0.001 | -0.46 | -0.28 |
| 14 | nA | 200 | 0.05 | 0.001 | -0.39 | -0.18 |
| 15 | nA | 200 | 0.05 | 0.001 | -0.41 | -0.17 |
| 16 | µA | 400 | 0.05 | 0.001 | -0.41 | -0.16 |

| | | | | | | |
|---|---|---|---|---|---|---|
| 17 | μA | 100 | 0.05 | 0.001 | -0.45 | -0.19 |
| 18 | nA | 100 | 0.05 | 0.001 | -0.41 | -0.18 |
| 19 | nA | 50 | 0.05 | 0.001 | -0.40 | -0.21 |
| 20 | nA | 200 | 0.05 | 0.001 | -0.42 | -0.16 |
| 21 | μA | 200 | 0.05 | 0.001 | -0.40 | -0.26 |
| 22 | μA | 200 | 0.05 | 0.001 | -0.40 | -0.29 |
| 23 | nA | 200 | 0.05 | 0.001 | -0.40 | -0.21 |
| 24 | μA | 200 | 0.05 | 0.001 | -0.40 | -0.20 |
| 25 | μA | 200 | 0.05 | 0.001 | -0.40 | -0.20 |
| 26 | μA | 200 | 0.05 | 0.001 | -0.40 | -0.48 |
| 27 | μA | 400 | 0.05 | 0.001 | -0.40 | -0.25 |
| 28 | μA | 200 | 0.05 | 0.001 | -0.40 | -0.31 |
| 29 | μA | 200 | 0.05 | 0.001 | -0.40 | -0.19 |
| 30 | μA | 200 | 0.05 | 0.001 | -0.40 | -0.59 |
| 31 | nA | 200 | 0.05 | 0.001 | -0.40 | -0.18 |
| 32 | nA | 200 | 0.05 | 0.001 | -0.40 | -0.17 |
| 33 | nA | 200 | 0.05 | 0.001 | -0.40 | -0.19 |
| 34 | μA | 100 | 0.05 | 0.001 | -0.40 | -0.24 |
| 35 | μA | 200 | 0.05 | 0.001 | -0.40 | -0.35 |
| 36 | μA | 200 | 0.05 | 0.001 | -0.40 | -0.11 |
| 37 | nA | 200 | 0.05 | 0.001 | -0.40 | -0.20 |
| 38 | nA | 200 | 0.05 | 0.001 | -0.40 | -0.20 |
| 39 | nA | 200 | 0.05 | 0.001 | -0.40 | -0.20 |
| 40 | nA | 200 | 0.05 | 0.001 | -0.40 | -0.20 |

**Table S1.** The detailed information of 40 folders selected from various experiments results. These folders have different current levels (μA and nA), frequencies (50 Hz, 100 Hz, 200 Hz and 400 Hz) and change point locations but share the same amplitude (0.05V) and potential step (0.001V).

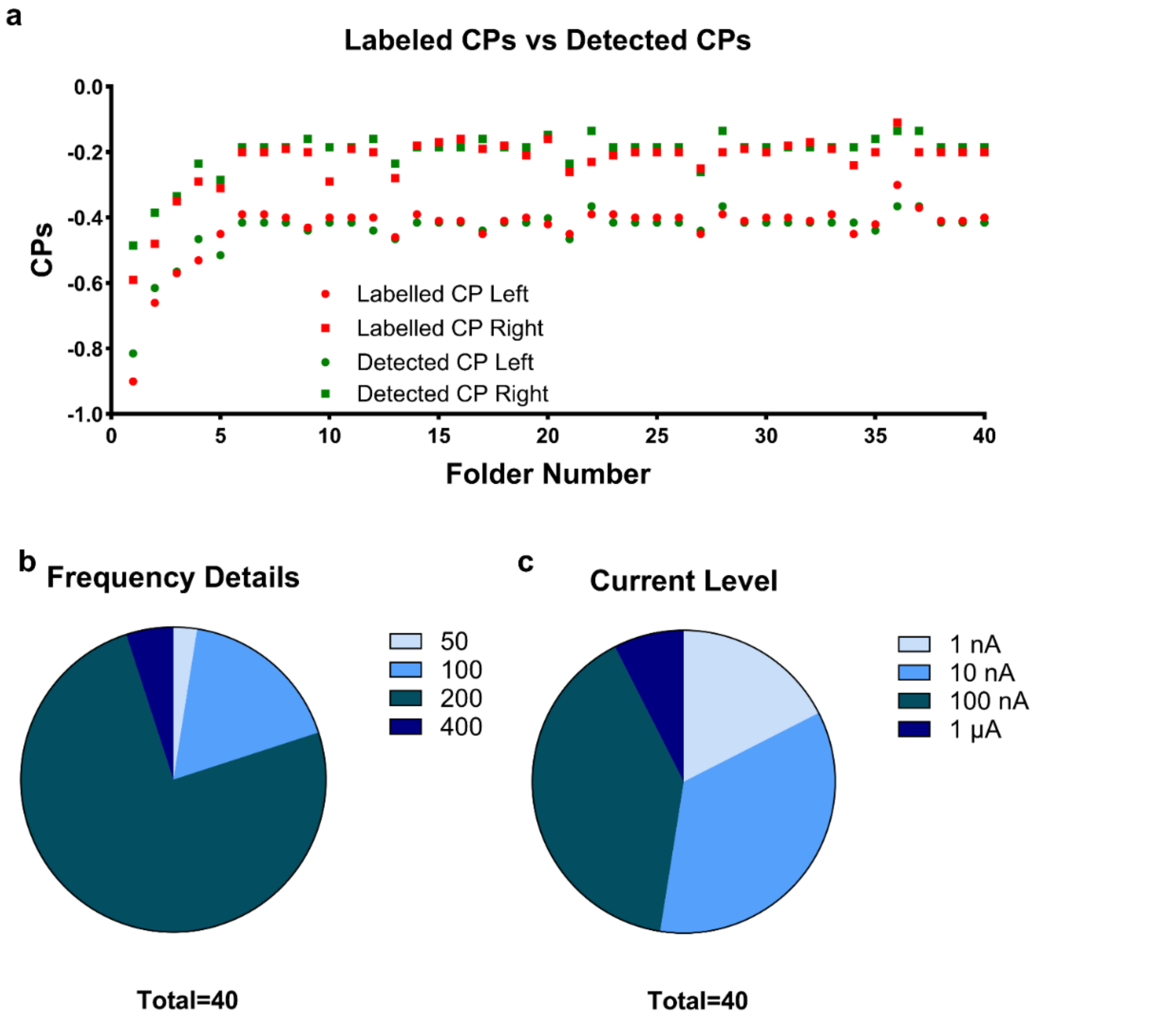

**Figure S2. a** Dataset details a human labeled CPs and detected CPs. The detected CPs share the similar locations with human labelled CPs on both left and right sides, which shows the reliability of CPD algorithm. **b** SWV measurement frequency details: 30 out of 40 folders have a frequency of 200 Hz, 7 folders have 100 Hz, 2 folders have 400 Hz, and one folder has 50 Hz. **c** SWV measurement current level details. Most folders have a current level between 10 nA and 1 μA while only 3 folders have a current level larger than 1 μA.

#### 1.4 CPD Performance Evaluation

To evaluate algorithm performance, the two detected CPs from each signal were compared against manually annotated CPs provided by researchers, ensuring consistency with human judgment. The algorithm-derived CPs were considered consistent with manually annotated CPs if the difference between them was within 10% of the interval separating the two CPs. Then we used the following scores to evaluate the peak consistency performance:

$$Scores = \frac{N_{consistent\ curves}}{N_{total\ curves}} \qquad (1)$$

, where $N_{total\ curves}$ is the 2,000 curves in dataset. When the score is over 0.8, the detected change points are considered consistent with the manually labeled ones. The details of scores for the 24 algorithms as a function of threshold is shown in **Figure S3**. Even when we extend the CPs searching range up to 80%, only BottomUp-rank and Dynp-rank can have a score over 0.8.

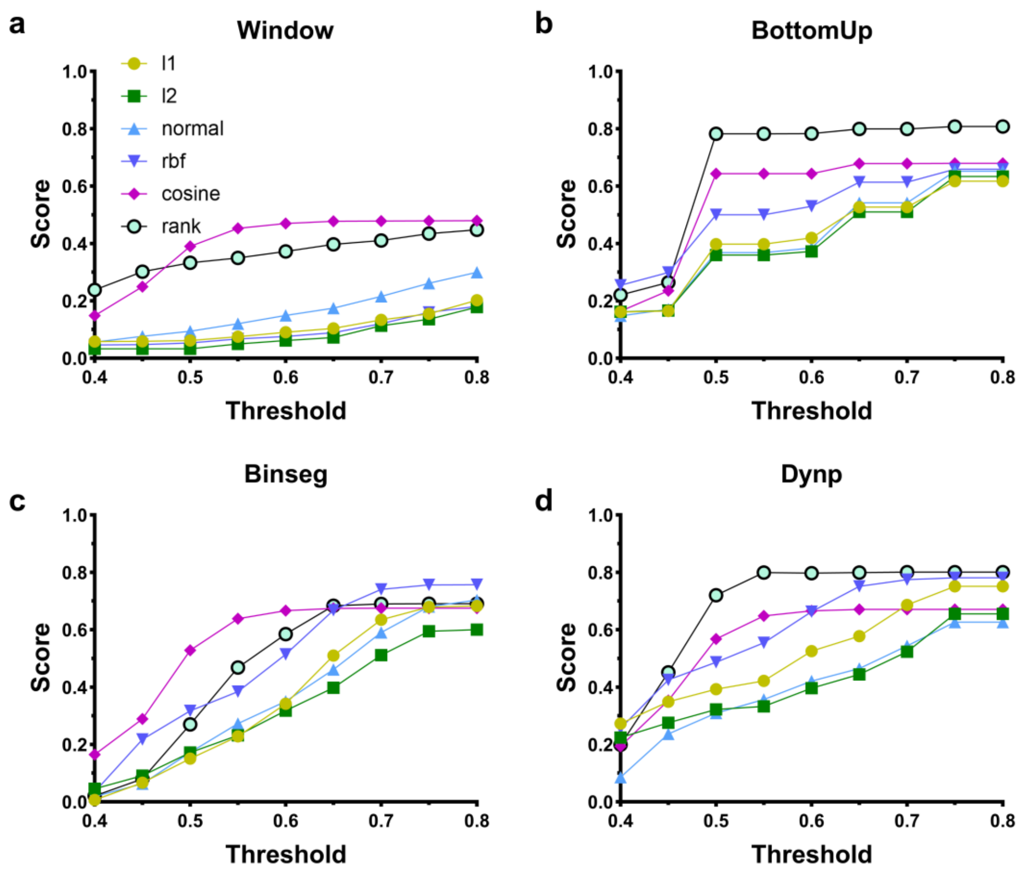

**Figure S3.** Scores of each CPD algorithm set after comparing the CPs from CPD and human labeling. BottomUp with rank and Dynp with rank have comparable scores that markedly surpass those of all other CPD algorithms. And both reach plateau phase after 65% threshold, which is consistent with the peak region size within the potential range of measurements.

### 2 Supplementary Note 2—Baseline fitting and screening

#### 2.1 Introduction to *pybaselines*

*pybaselines* is a Python library[10], designed for the baseline fitting of experimental data. Among all the included baseline fitting algorithms, we selected 30 algorithms for baseline fitting. Details of these algorithms are listed in **Table S2**.

#### 2.2 Details of Baseline Shape Check

The threshold for baseline over raw data was set as 10% in the peak region. Meanwhile, a baseline is considered invalid if the area enclosed by the baseline and the CP connection line exceeds 30% of the area enclosed by the CP connection line and the original curve.

| Type | Algorithms |
| --- | --- |
| Polynomial Baselines (Up to order 3) | Goldindec (Goldindec Method), imodpoly (Improved Modified Polynomial), modpoly (Modified Polynomial), poly (Regular Polynomial), quant_reg (Quantile Regression),penalized_poly (Penalized Polynomial) |
| Whittaker Baselines | Airpls (Adaptive Iteratively Reweighted Penalized Least Squares), arpls (Asymmetrically Reweighted Penalized Least Squares), aspls (Adaptive Smoothness Penalized Least Squares), derpsalsa (Derivative Peak-Screening Asymmetric Least Squares Algorithm), drpls (Doubly Reweighted Penalized Least Squares), iarpls (Improved Asymmetrically Reweighted Penalized Least Squares), psalsa (Peaked Signal's Asymmetric Least Squares Algorithm) |
| Penalized Spline Version Baselines | irsqr (Iterative Reweighted Spline Quantile Regression), mixture_model (Mixture Model), pspline_airpls (Penalized Spline Version of airpls), pspline_arpls (Penalized Spline Version of arpls), pspline_aspls (Penalized Spline Version of aspls), pspline_derpsalsa (Penalized Spline Version of derpsalsa), pspline_drpls (Penalized Spline Version of drpls), pspline_iarpls (Penalized Spline Version of iarpls), pspline_mpls (Penalized Spline Version of mpls), pspline_psalsa (Penalized Spline Version of psalsa) |

| | |
|---|---|
| Classification Baselines | cwt_br (Continuous Wavelet Transform Baseline Recognition), dietrich (Dietrich's Classification Method), fabc (Fully Automatic Baseline Correction), fastchrom (FastChrom's Baseline Method), golotvin (Golotvin's Classification Method), rubberband (Rubberband Method), std_distribution (Standard Deviation Distribution) |

**Table S2.** Baseline fitting algorithms from *pybaselines* package, including 4 types of fitted baseline: polynomial baselines, Whittaker baselines, penalized spline version baselines and classification baselines

#### 2.3 Types of SWV curves

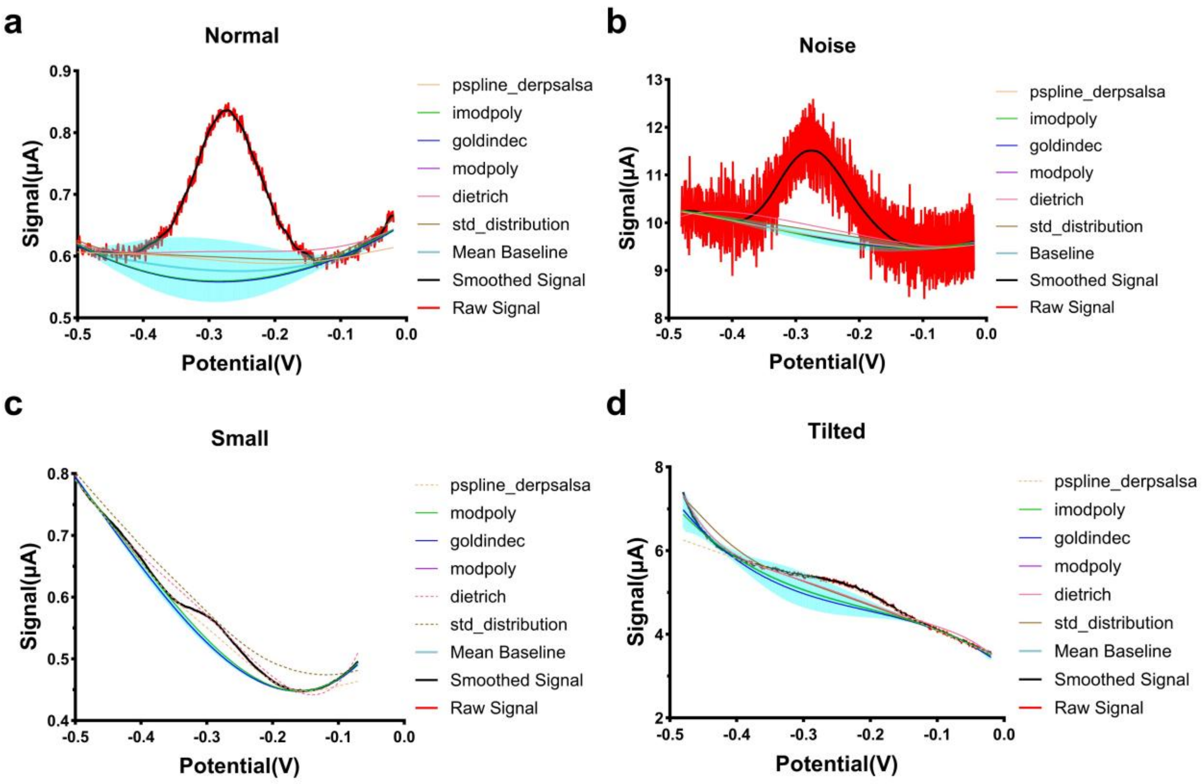

**Figure S4.** SWV curve types processed by A-PACE default algorithm set. **a** An example of a normal SWV curve. All baseline fitting algorithms in default algorithm set can pass the shape check, outlier check and the median value of passed baselines decide the final baseline output. **b** An example of a noise SWV curve: with level 3 smoothing, the noise curve is converted to normal curve and subsequently subjected to analogous processing by A-PACE. **c** An example of a small SWV curve: Owing to algorithmic redundancy, several algorithms passed the shape-check stage even when others did not, ultimately yielding an appropriate baseline. **d** An example of a tilted SWV curve: Although most baselines passed the shape check, a subset of curves still differed markedly from the others and was subsequently discarded during outlier detection.

#### 2.4 Algorithm Set Selection

For each signal curve, 30 algorithms in **Table S2** were applied for baseline fitting. And each baseline was screened with the shape check in **Section 2.2**. Among 2,000 curves in the dataset, the success rate (SR) can be statistically determined:

$$Success\ Rate = 1 - \frac{N_{dropout}}{N_{total}} \quad (2)$$

Then the peak heights were obtained from the passed baselines and raw signals and composed a time-series data set for each folder. Then such time-series data were further smoothed with Locally Weighted Scatterplot Smoothing (LOESS), with a window size of 0.3 and 3 iterations. And NMSE (Normalized Mean Square Error) can be calculated between the raw time-series data and the Loss smoothed results:

$$NMSE = \frac{\sum_{i=1}^{N_{folder}} \sum_{t=1}^{N_{curves}} [\,(I_{i,t,A-PACE} - I_{i,t,Loss})^2/(I_{i,max} - I_{i,min})^2]}{N_{folder}} \quad (3)$$

, where $N_{Folder}$ is the total number of time-series data folders while $N_{curves}$ is the curves in each folder. $I_{i,max}$ and $I_{i,min}$ are the maximum and minimum signal value of smoothed signals within each folder.

Then for baselines derived from the same CPD algorithm, we exhaustively combined them, resulting in 2*1023 possible combinations. For each combination, all baselines were subjected to a shape verification procedure. If more than four baselines remained after this step, an extreme value detection analysis was subsequently conducted. The local outlier factor outlier detection was carried out with scikit-learn package[11]. The number of neighbors to consider when calculating the local density of a data point 5 while the proportion of outliers in the dataset is 'auto'.

The following **Table S5** further provides more insight of the A-PACE default algorithm set. With all the baseline fitting algorithm set with BottomUp for CPD and 100% SR, the *pspline_derpsalsa*, *goldindec* and *modpoly* occur most frequently in the top algorithms, which means that these three algorithms can provide stable baseline fitting results with high SR. The algorithm set of these three algorithms has a SR of 100% (0.26 for $Z_{SR}$) and NMSE of 0.0200 (-1.86 for $Z_{NMSE}$), which are very close to default algorithm set. As integral components of the default algorithm (**Table S4**), these three algorithms can be considered to play a dominant role in determining its performance.

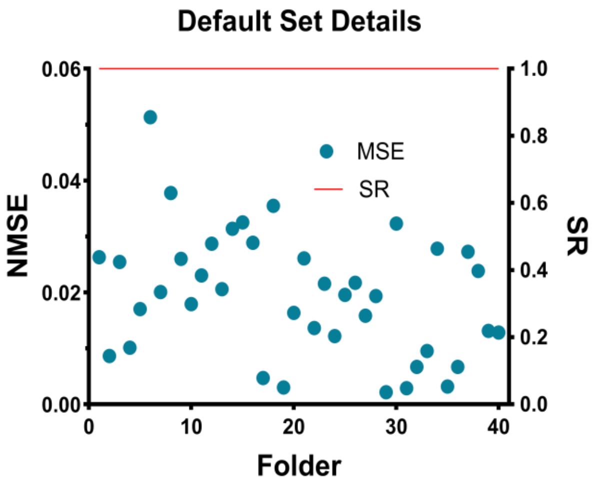

**Figure S5.** Performance of default baseline fitting algorithm set within each folder. The default algorithm has a SR of 100% over all 40 folders and in most folders the NMSE is under 0.04, which indicates excellent stability across the time-series.

| CPD Algorithms | Baseline Fitting Algorithms |
|---|---|
| Dynp-rank | quant_reg, derpsalsa, pspline_airpls, pspline_mpls, pspline_derpsalsa, dietrich, pspline_psalsa, psalsa, std_distribution, fastchrom |
| Bottom-rank | quant_reg, pspline_derpsalsa, derpsalsa, imodpoly, penalized_poly, goldindec, modpoly, dietrich, std_distribution, fastchrom |

**Table S3.** Baseline fitting algorithms as 1023 combinations for each CPD algorithm.

| CPD Algorithm Set | Baseline Fitting Algorithms |
|---|---|
| BottomUp + rank | pspline_derpsalsa, imodpoly, goldindec, modpoly, dietrich, std_distribution |

**Table S4.** CPD algorithm and baseline fitting algorithms details of the A-PACE default algorithm set.

| | Number (%) in Top 10 | Number (%) in Top 20 | Number (%) in Top 100 |
|---|---|---|---|
| quant_reg | 0 (0) | 3 (15) | 43 (43) |
| pspline_derpsalsa | 10 (100) | 18 (90) | 77 (77) |
| derpsalsa | 0 (0) | 0 (0) | 13 (13) |
| imodpoly | 7 (70) | 13 (65) | 78 (78) |
| penalized_poly | 3 (30) | 6 (30) | 68 (68) |
| goldindec | 9 (90) | 19 (95) | 80 (80) |
| modpoly | 9 (90) | 19 (95) | 81 (81) |
| dietrich | 4 (40) | 6 (30) | 22 (22) |
| std_distribution | 7 (70) | 12 (60) | 50 (50) |
| fastchrom | 2 (20) | 6 (30) | 42 (42) |

**Table S5.** Single baseline fitting algorithm performance in top 10, 20, 100 algorithm set, respectively. *pspline_derpsalsa* is present in 100 % of the top 10 algorithm sets, whereas *goldindec* and *modpoly* each appear in 90 %. These three baseline fitting algorithms far outperform the rest and continue to dominate within the top 20 and top 30 algorithm sets.

### 3 Supplementary Note 3—Data interaction and visualization workflow

#### 3.1 Rationale to design GUI and key advantages

The graphical user interface (GUI) was meticulously designed to optimize data processing workflow for researchers and laboratory technicians, offering an accessible and efficient platform for real-time analysis and post-experiment processing. The system facilitates automated file extraction and organization through a locally executed Tkinter GUI, enabling seamless interaction without requiring advanced technical expertise. Users can apply pre-trained algorithms, upload custom models, and dynamically adjust parameters such as peak height, frequency thresholds, and concentration values through intuitive sliders and dropdown menus. Advanced visualization capabilities include both 2D and 3D graph generation, with customizable axes, overlays, and color schemes, empowering researchers to derive deeper insights into complex datasets. Real-time graph updates further enhance usability, particularly for time-series data analysis.

The architecture employs a Flask-based backend, integrating robust libraries such as numpy, scipy, and matplotlib for computational processing. For interactive 3D visualizations, the GUI leverages Plotly, providing high-performance, user-friendly rendering of complex datasets. Error handling mechanisms are built into the system, including automatic file validation to ensure compatibility, real-time progress indicators during data processing, and detailed error messages for rapid troubleshooting. User support is further reinforced through an interactive help section and descriptive tooltips, making the system accessible to both novice and experienced users.

Distinguishing features of the GUI include batch processing for large-scale data workflow, the ability to dynamically integrate custom algorithms, and export options for processed data and visualizations in formats such as .csv, .png, and .pdf. Additionally, the system maintains a comprehensive history of processed files and results, allowing users to revisit prior analyses for comparison or further refinement. Designed with a user-centric approach, the interface emphasizes clarity and accessibility, employing a clean, minimalist layout with logically grouped controls to streamline workflows and reduce cognitive load. Accessibility features such as keyboard shortcuts and screen reader support further enhance usability across diverse user needs.

This GUI represents a significant advancement in laboratory data analysis, offering a powerful, flexible tool for efficient and accurate processing. Future developments could include integration with cloud-based storage and processing platforms, collaborative tools for multi-user engagement, and direct support for training and evaluating machine learning models within the interface. By uniting automation, customization, and advanced visualization in a single platform, this GUI sets a new standard for data analysis in experimental and computational research.

#### 3.2 Key features of our GUI

For post-data processing mode, with the data extracted from *.pssession* file (from MultiTrace) or *.csv* file, the tool will utilize all CPU cores for CPD and baseline fitting/screening. Then the final results will be divided into 2 groups: ’Pass’ and ’Fail’. For ’Pass’ group, signals with fitting baselines and peak value were shown for user review (**Figure S6a**). Meanwhile, users can efficiently filter results in batches and compare curves with a overlay graph. (User Manual of A-PACE - Post-Data Processing Pass). For ’Fail’ group, the raw data of signals are shown. For signals that should have succeeded, one of the most common causes of failure is incorrect CP positioning, possibly due to an unsuitable peak region threshold (**Figure S6b**). Thus, manually inputting CPs can correct this error and allow curves with reasonable results to be moved back to the Pass group (**Figure S6c**). For all curves in the ‘Pass’ group, a data table summarize all the available information, including ‘curve path and index’, ‘Data and Time’, ‘Frequency (Hz)’, ‘Amplitude(V)’, ‘E Step(V)’, ‘peak Height($\mu$A)’, ‘Channel Number’, and ‘Concentration (mM)’. Meanwhile, batch adding or modifying relevant information in the table can be realized by specifying the index range. And these data can be used for both 2D and 3D plottings and will be saved in a JSON file for review.

A-PACE, developed with an HTML-based interface using Flask and the PalmSens SDK, supports data inputs from *.csv* and *.pssession* files. Users can select real-time detection or post-processing modes. Real-time detection processes files continuously, searching and updating results within seconds, with a sliding window algorithm addressing peak drift by averaging change point locations. In post-processing, all CPU cores are utilized for CPD and baseline fitting, categorizing results into passed or failed. Users can batch filter results, modify change points to improve quality, and access measurement details for sequence plotting.

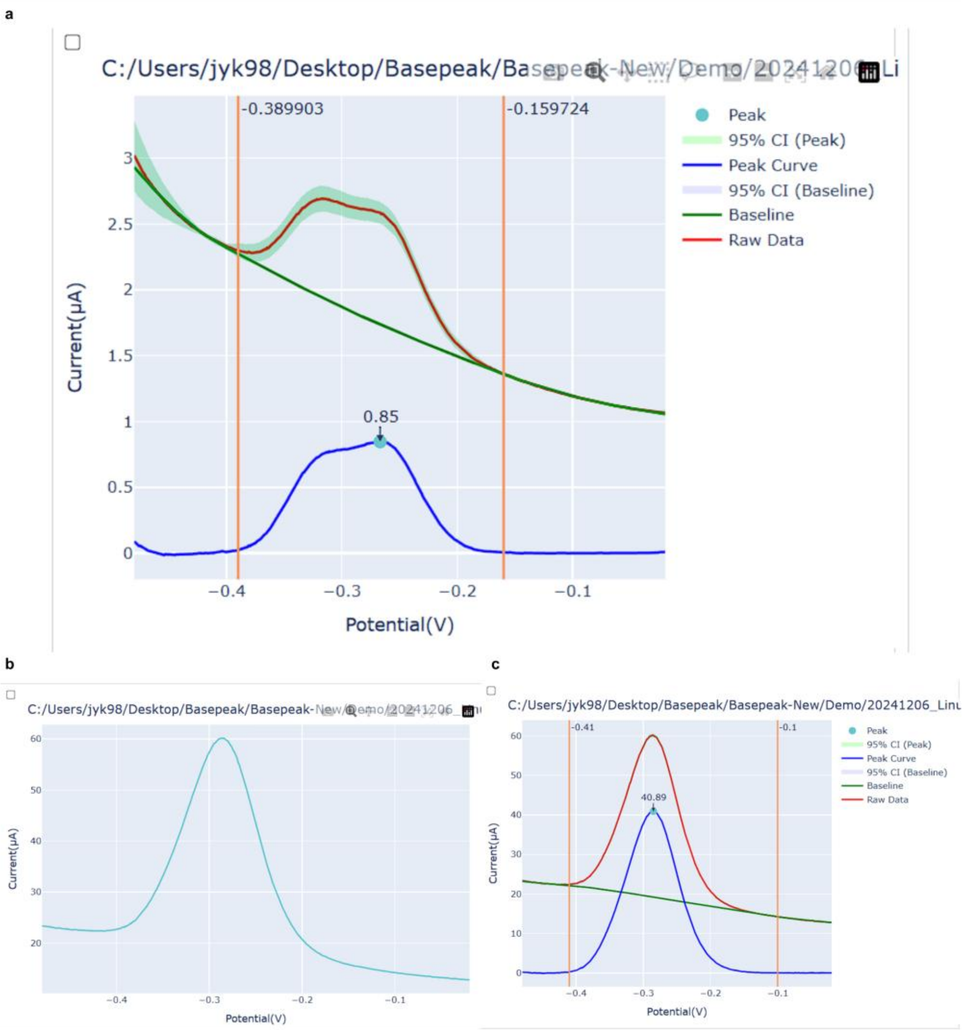

**Figure S6.** Snapshot of Output Groups. **a** An example of a curve in the pass group. Analysis details includes including raw data, change points, fitted baseline and net peak. **b** An example of a curve in the fail group. The lack of change point information means the failure is caused by improper peak region threshold input or CPD algorithm. **c** An example of an adjusted curve from the fail group. User can manually input the CPs of all the curve in the fail group and A-PACE can carry out the baseline fitting based on the new CPs for new analysis results.

### 4 Supplementary Note 4—Case Studies

As one of the most important subclasses of aminoglycoside antibiotics (AGs), kanamycin has been widely used for Gram-positive and Gram-negative bacteria infection treatment[12]. Since a high plasma level of AGs may cause ototoxicity and nephrotoxicity, serum levels have to be controlled within a narrow therapeutic range[13]. Compared with traditional AGs detection method, like liquid chromatography (LC), LC-mass spectrometry (MS), capillary electrophoresis-MS, microbial assays, and enzyme-linked immunoassays, electrochemical biosensors have been widely used for AGs detection due to its strong specificities and high sensitivities[14].

#### 4.1 Details of different case studies

A-PACE reliably analyzes extended-duration EAB signals, even for month-long *in vitro* measurements and week-long *in vivo* tests, providing stable baselines, well-defined change points, and accurate peak heights despite significant signal decline. Compared to default parameter settings, A-PACE with selected algorithm set demonstrates enhanced success rates, improved sensitivity to kanamycin concentration changes, and stable performance through long-term incubations and free-moving animal studies. Although minimal manual adjustments were needed at later time points, A-PACE's overall accuracy and efficiency make it a powerful tool for extended biosensor monitoring.

#### 4.2 Examples of *in vitro* serum data

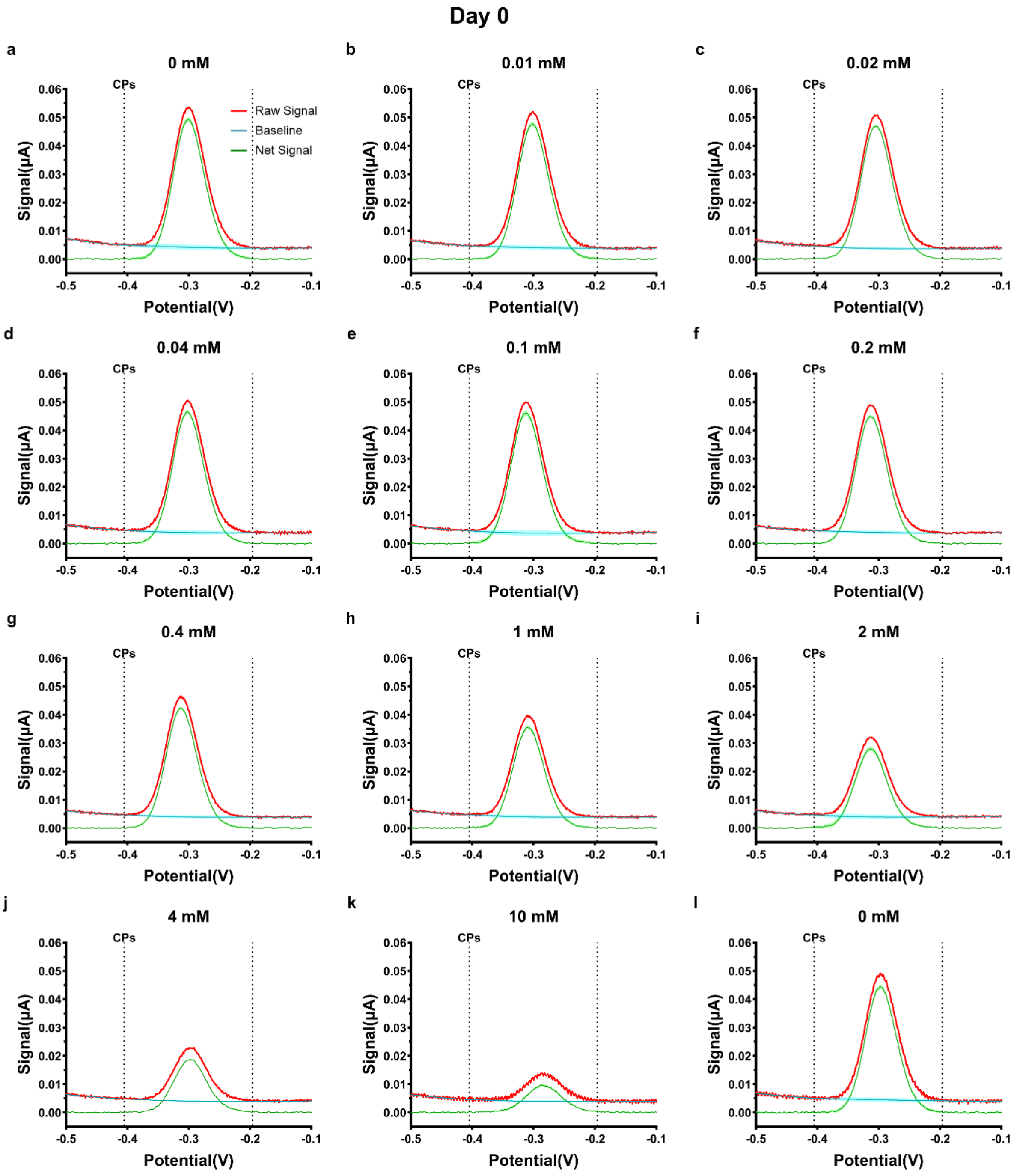

**Figure S7.** *in vitro* day 0 analysis details. The subfigures show the raw signals from measurements, related baselines from A-PACE default algorithm and net peak results over different concentrations of kanamycin ranging from 0.01–10 mM.

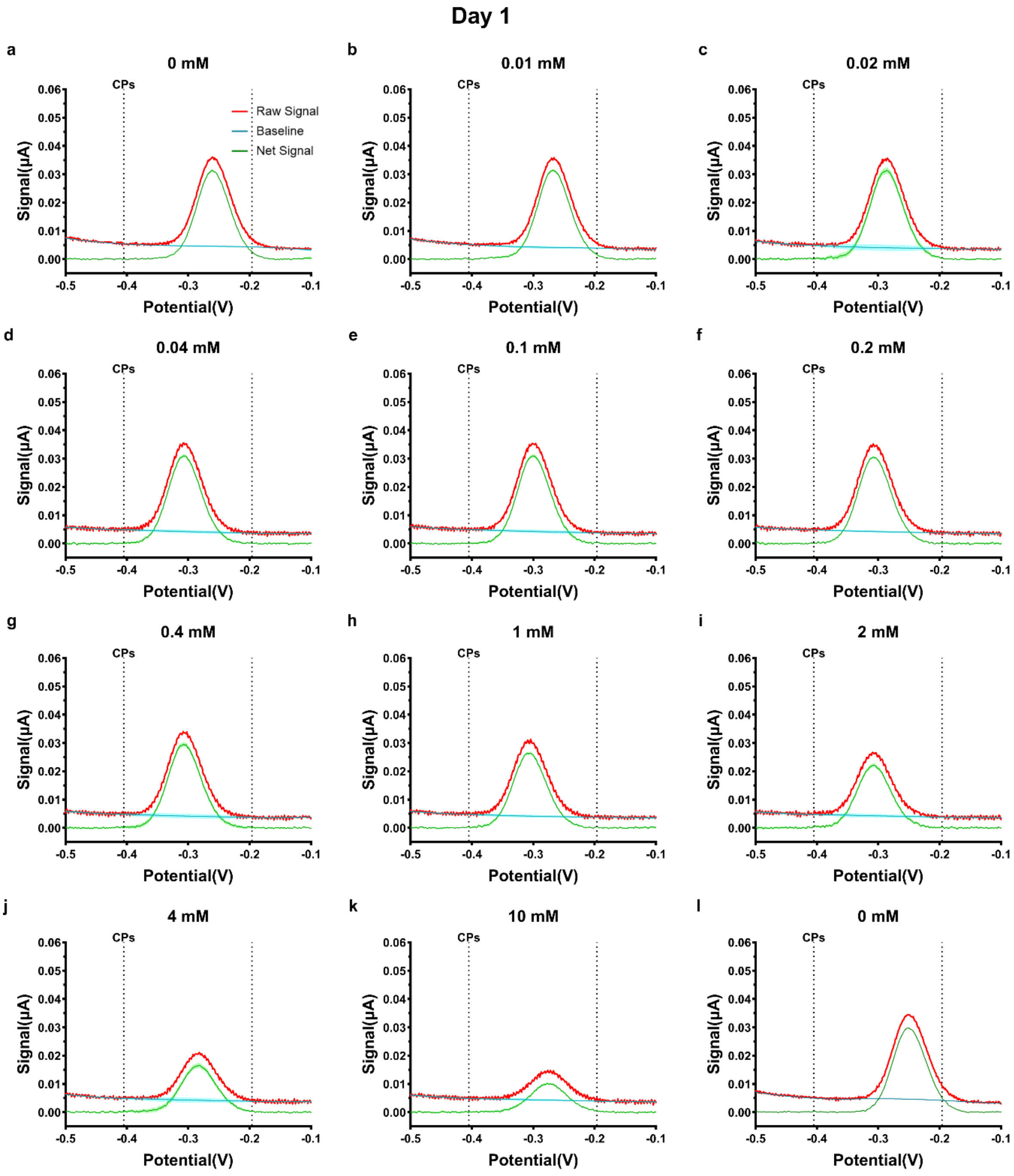

**Figure S8.** *in vitro* day 1 analysis details. Compared with signals on day 0, peak regions initially show a low amplitude on day 1.

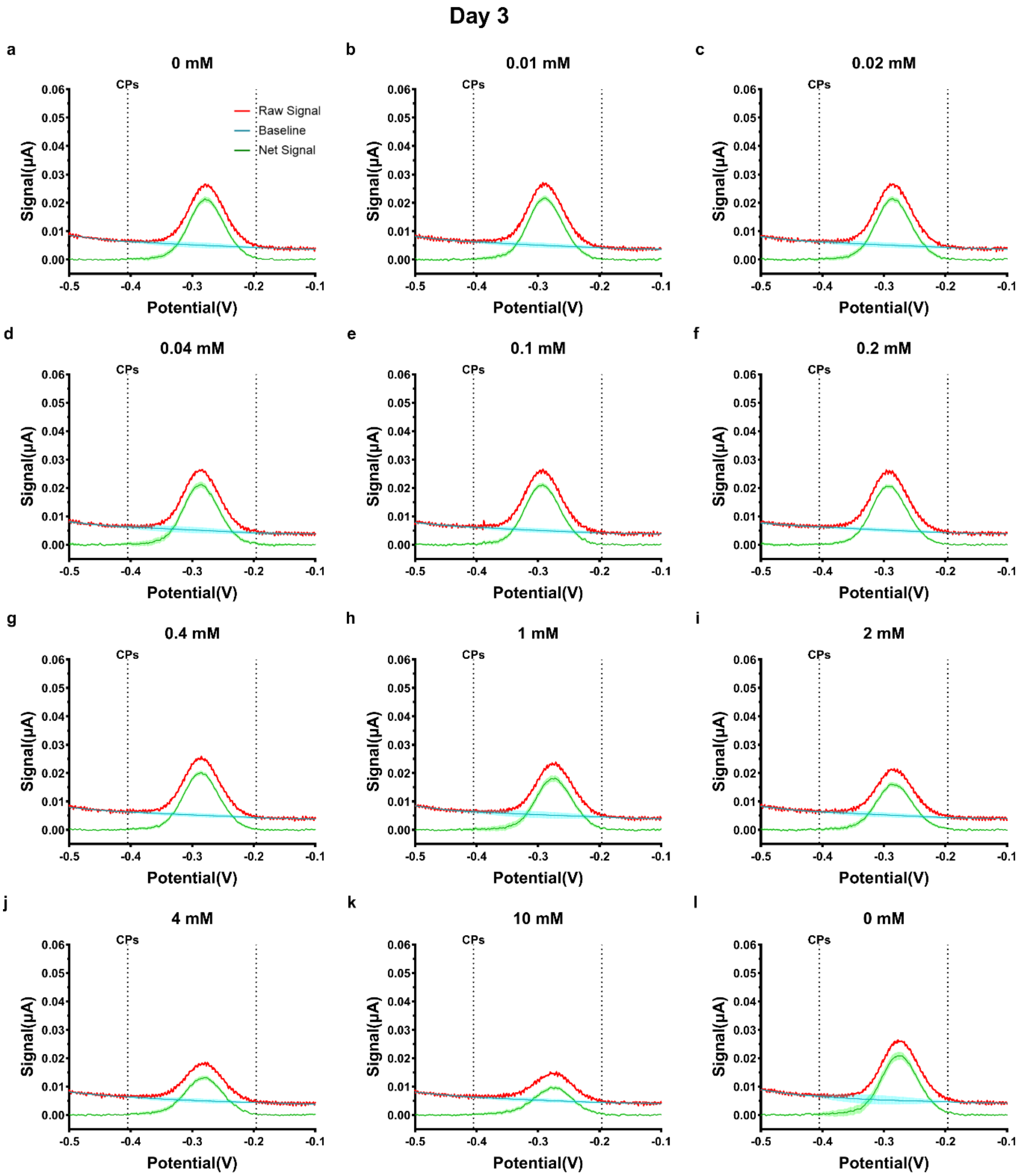

**Figure S9.***in vitro* day 3 analysis details.

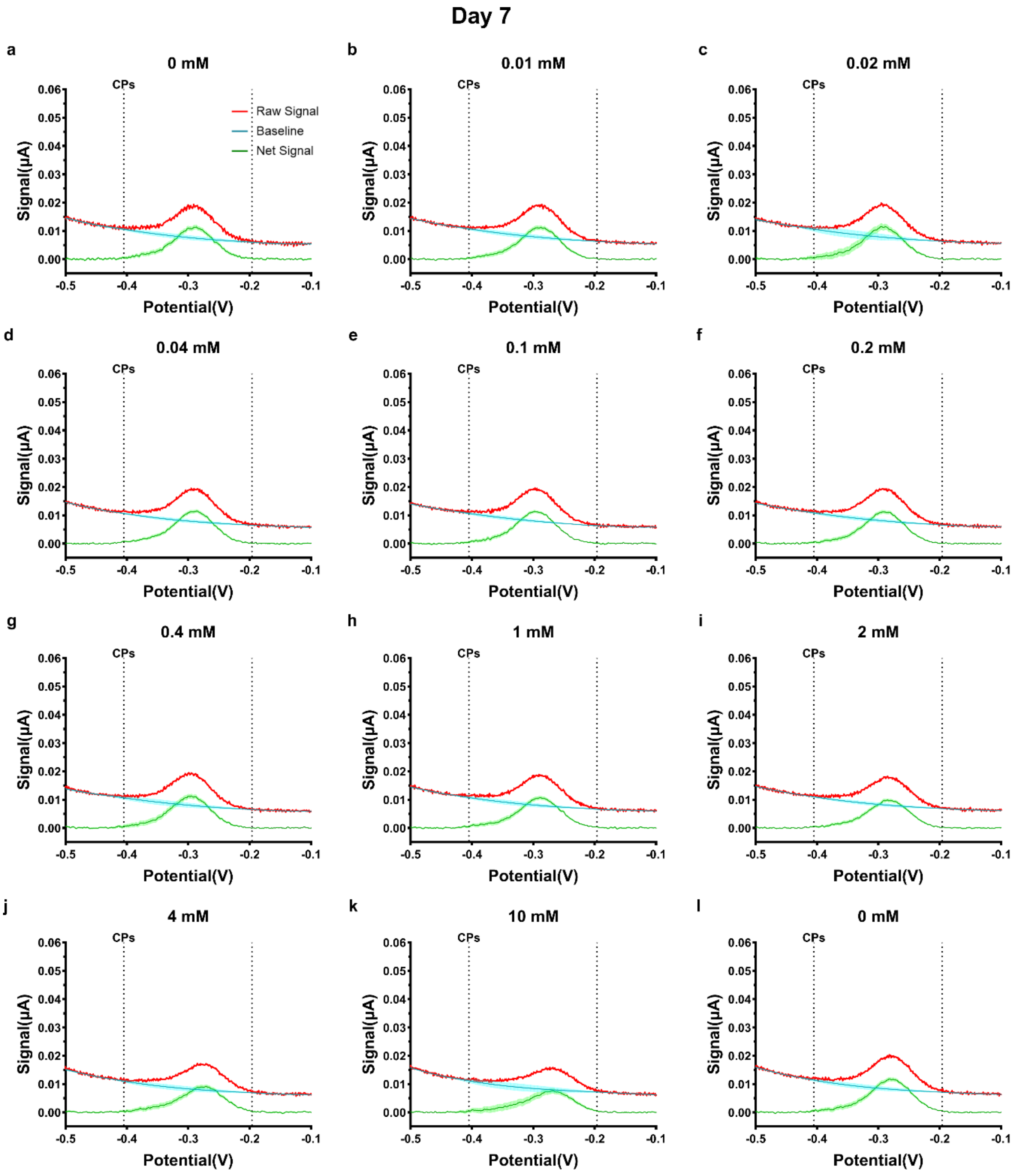

**Figure S10.** *in vitro* day 7 analysis details. Signals start to become tilted for all concentrations.

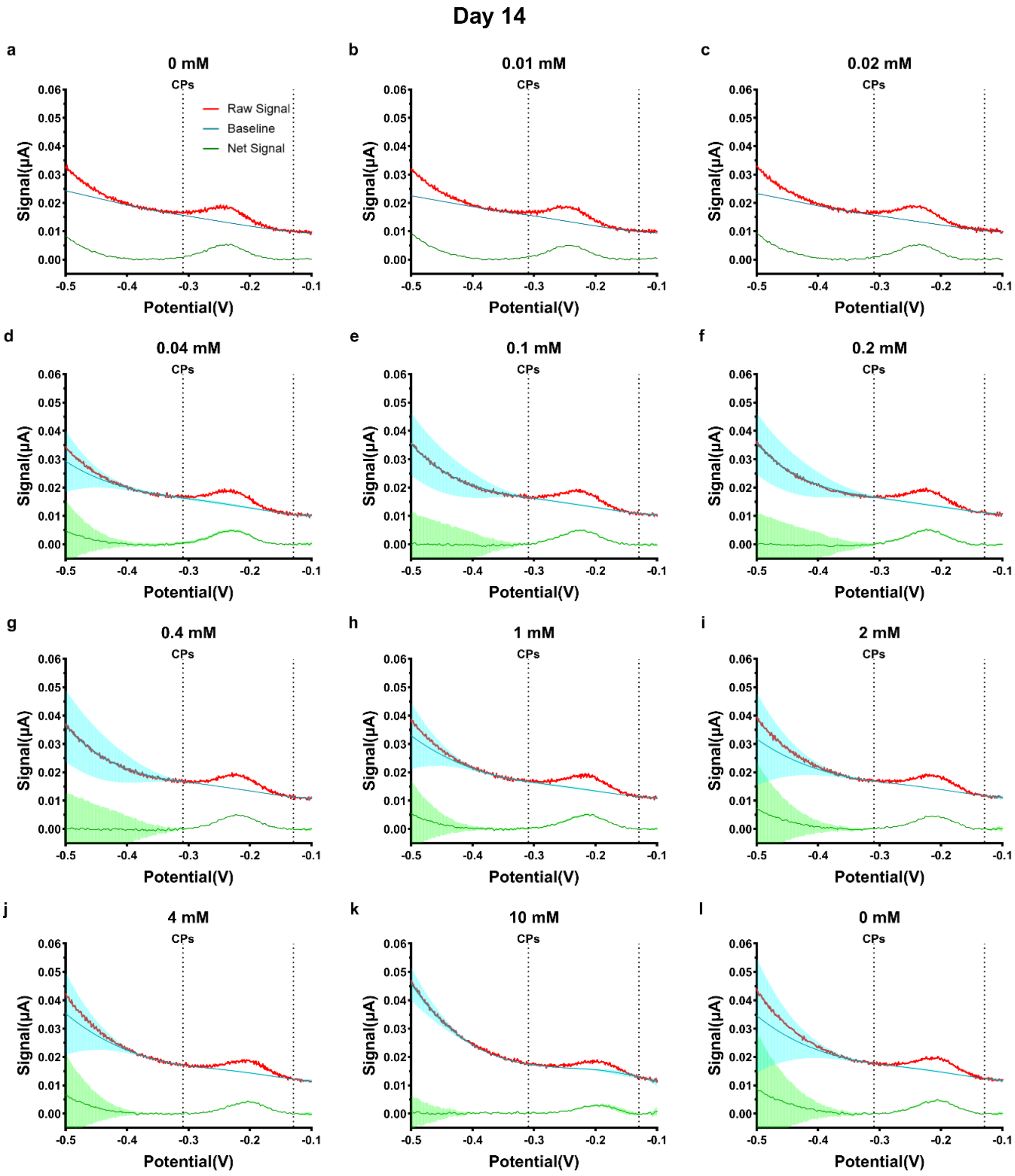

**Figure S11.** *in vitro* day 14 analysis details. More tilted and small peak signals are obtained on day 14.

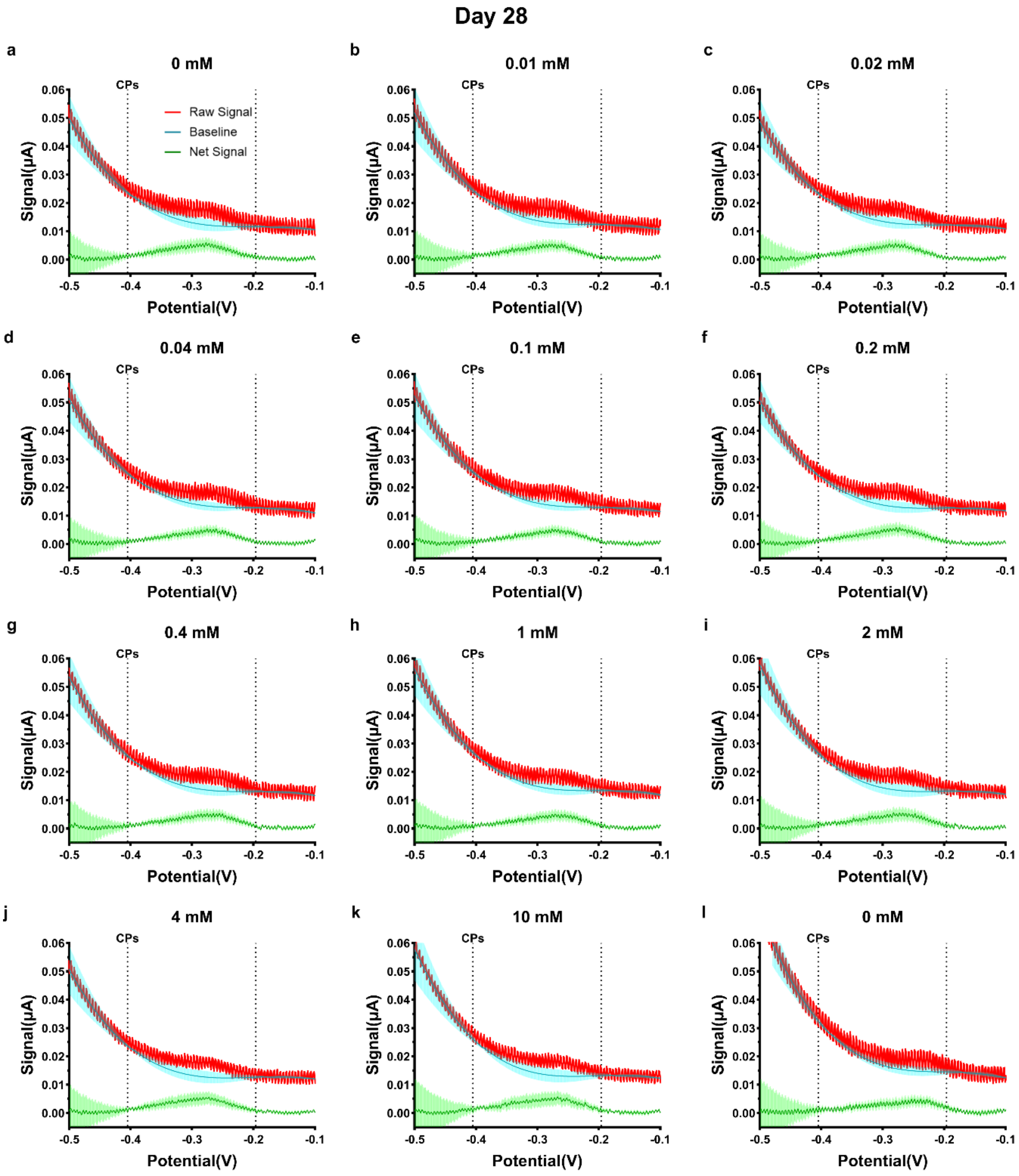

**Figure S12.** *in vitro* day 28 analysis details. The signals on day 28 are highly tilted and has high SNR, which can hardly be processed by manual labelling.

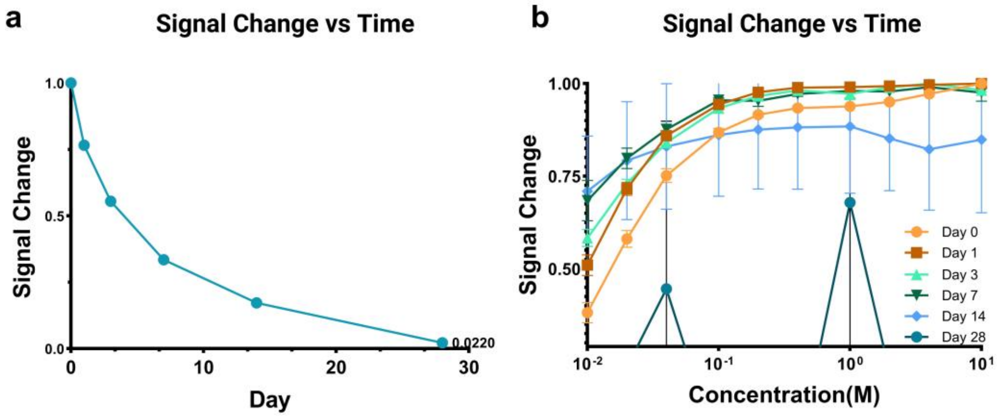

**Figure S13. a** 28 days' *in vitro* signal degradation with time (from MultiTrace). **b** *in vitro* signal degradation with concentration over 28 days (from MultiTrace).

#### 4.3 Examples of *in vivo* blood data

To address the challenge of differentiating target-induced signal gain from drift *in vivo*, the stability was assessed by calibrating sensors in Phosphate-buffered saline (PBS) with defined kanamycin concentrations before and after intravenous implantation. After implantation and recovery, the rats moved freely, subjecting the electrode probes to continuous blood flow and mechanical strain.

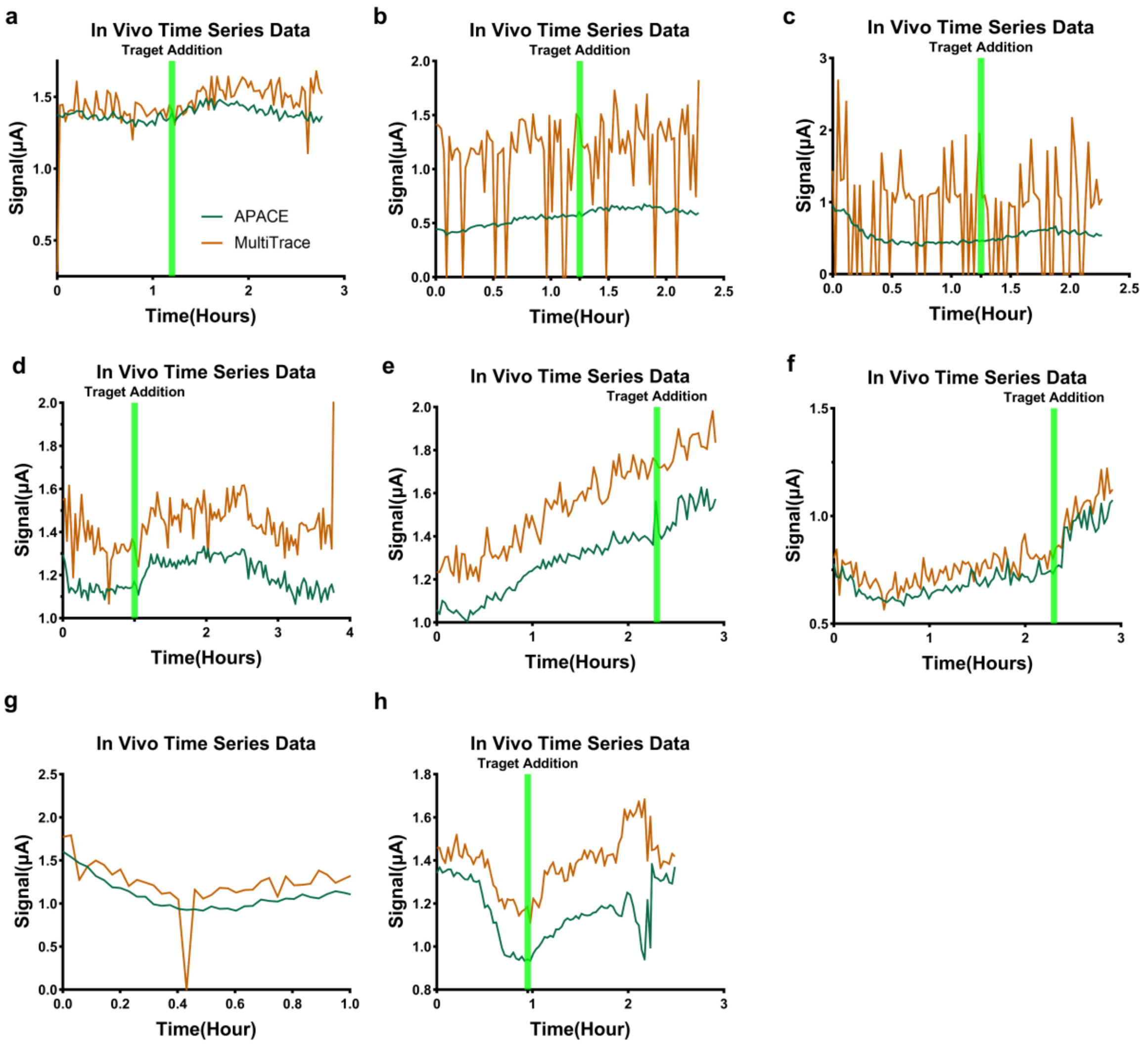

**Figure S14.** *in vivo* time-series signal processing results compared with those from MultiTrace. MultiTrace have failure analysis in 3 out of 8 cases), but A-PACE has a SR of 100% with much smoother results.

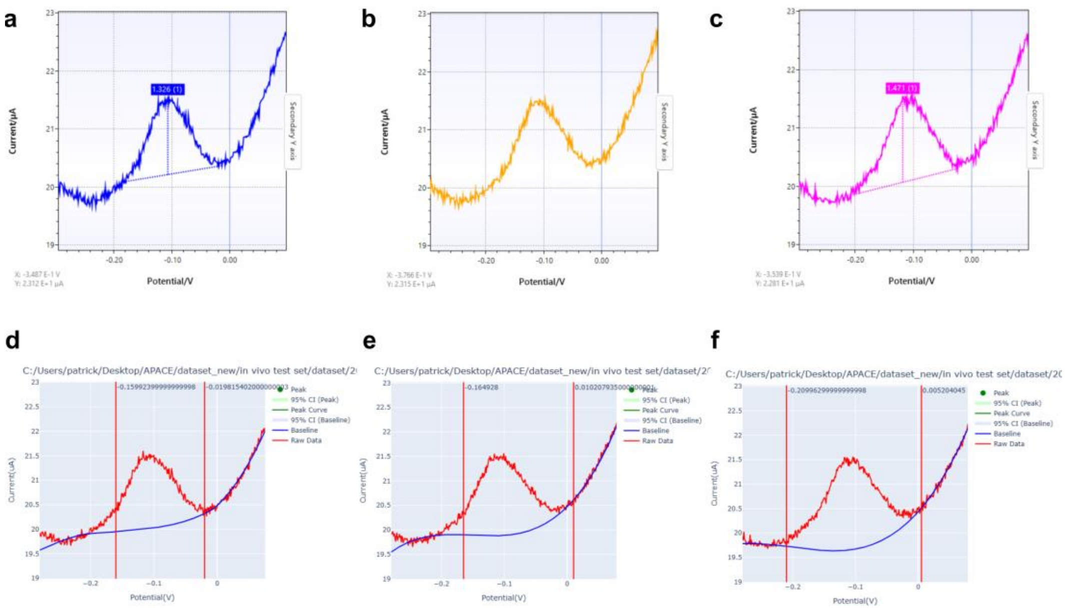

**Figure S15. a-c** MultiTrace results from three consecutive SWV measurements. MultiTrace fails on the middle one. **a** before failure time point. **b** at failure time point. **c** after failure time point. **d-f** A-PACE find similar baselines over these three cases. **d** before failure time point. **e** at failure time point. **f** after failure time point.

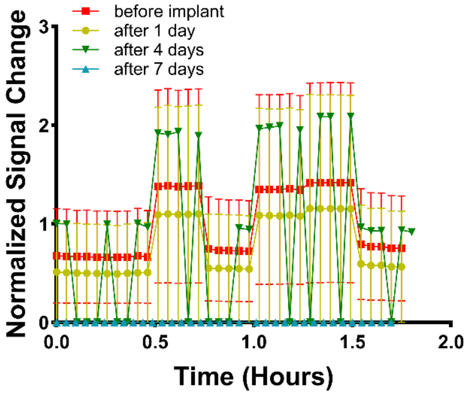

**Figure S16.** *in vivo* signal variation from MultiTrace before and after sensor implantation into the animal for 1, 4, 7 days, respectively. The results from before implant and after day 1 have response to different concentrations. After day 4, MultiTrace starts failing in some cases while all cases fail after day 7. All results have larger error bars, showing unstable analysis results over week-long continuous electrochemical measurements.

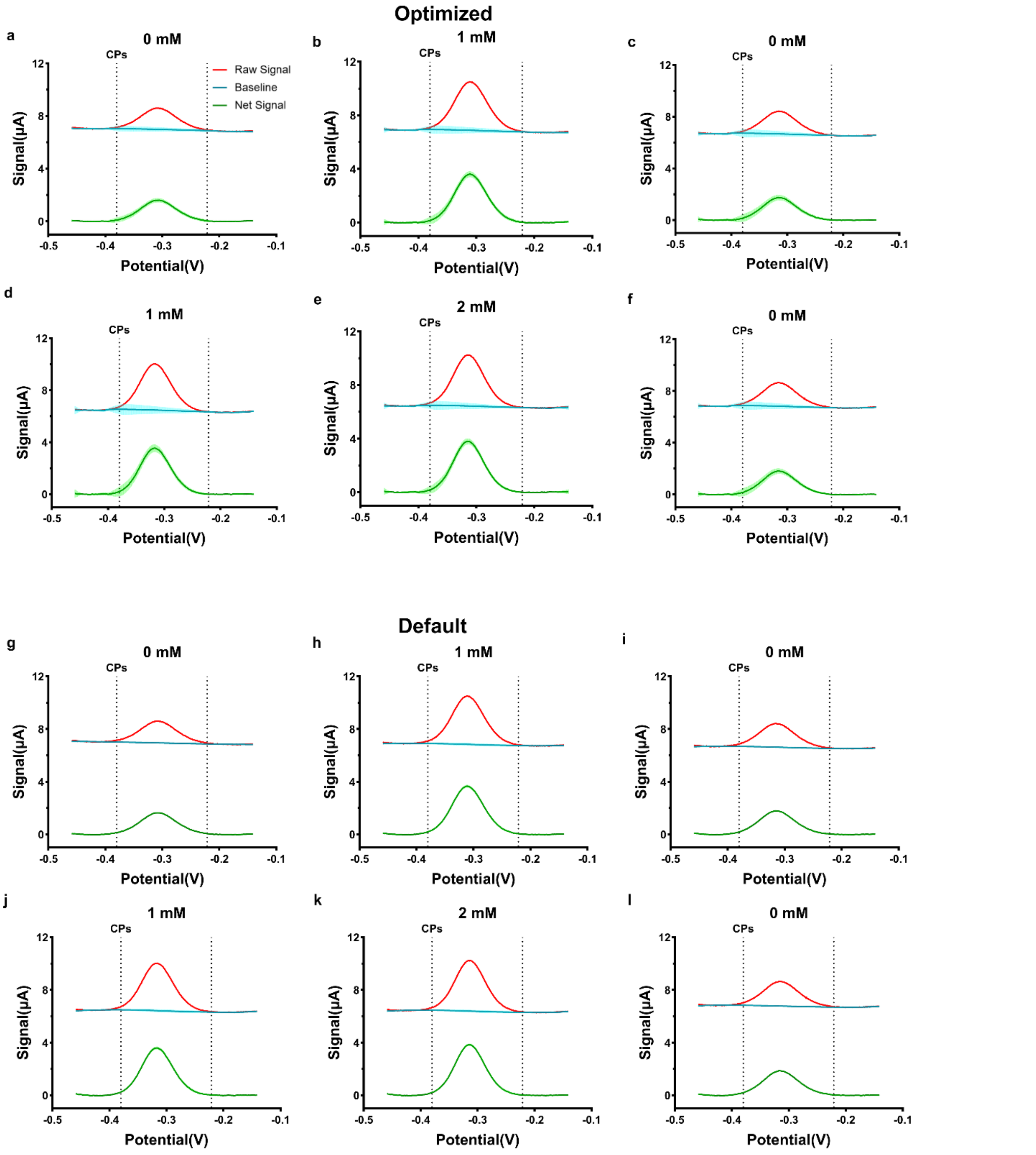

**Figure S17.** *in vivo* before implant fitting details with default algorithm set and optimized algorithm set. The subfigures show the raw signals from measurements, related baselines from A-PACE default algorithm and net peak results over different concentrations before implant.

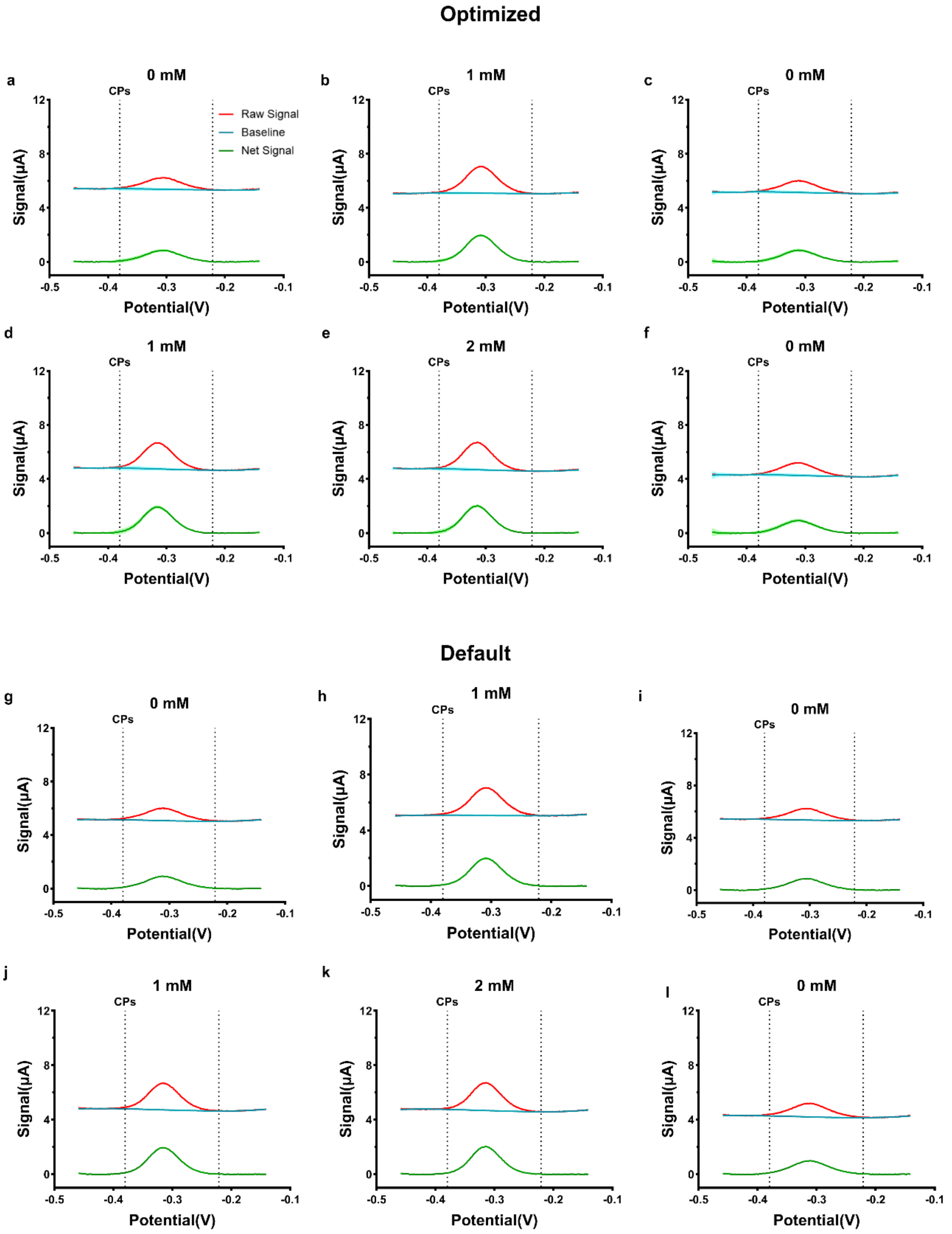

**Figure S18.** *in vivo* day 1 fitting details with default algorithm set and optimized algorithm set. Compared with signals before implant, peak regions initially show a low amplitude on day 1.

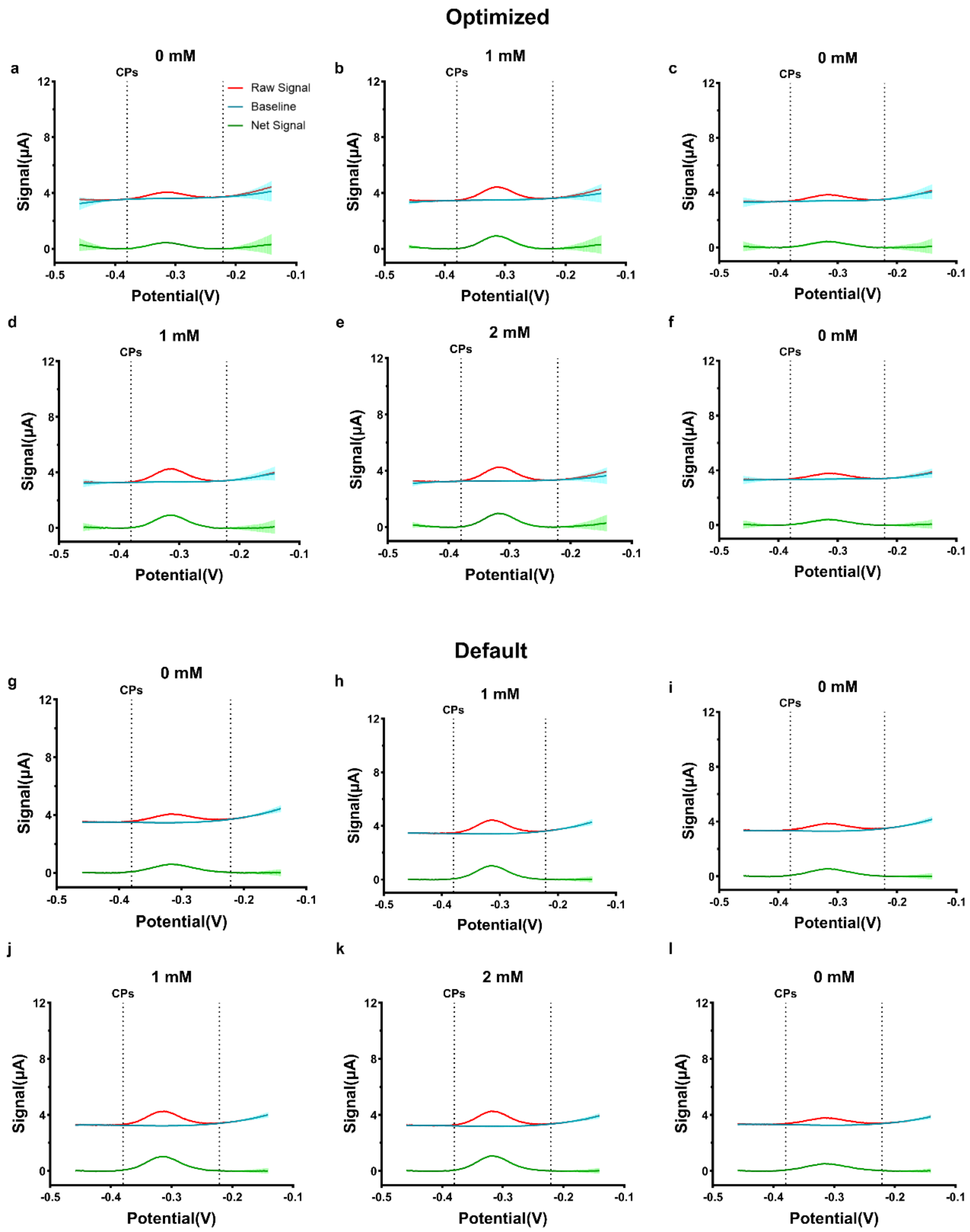

**Figure S19.** *in vivo* day 4 fitting details with default algorithm set and optimized algorithm set.

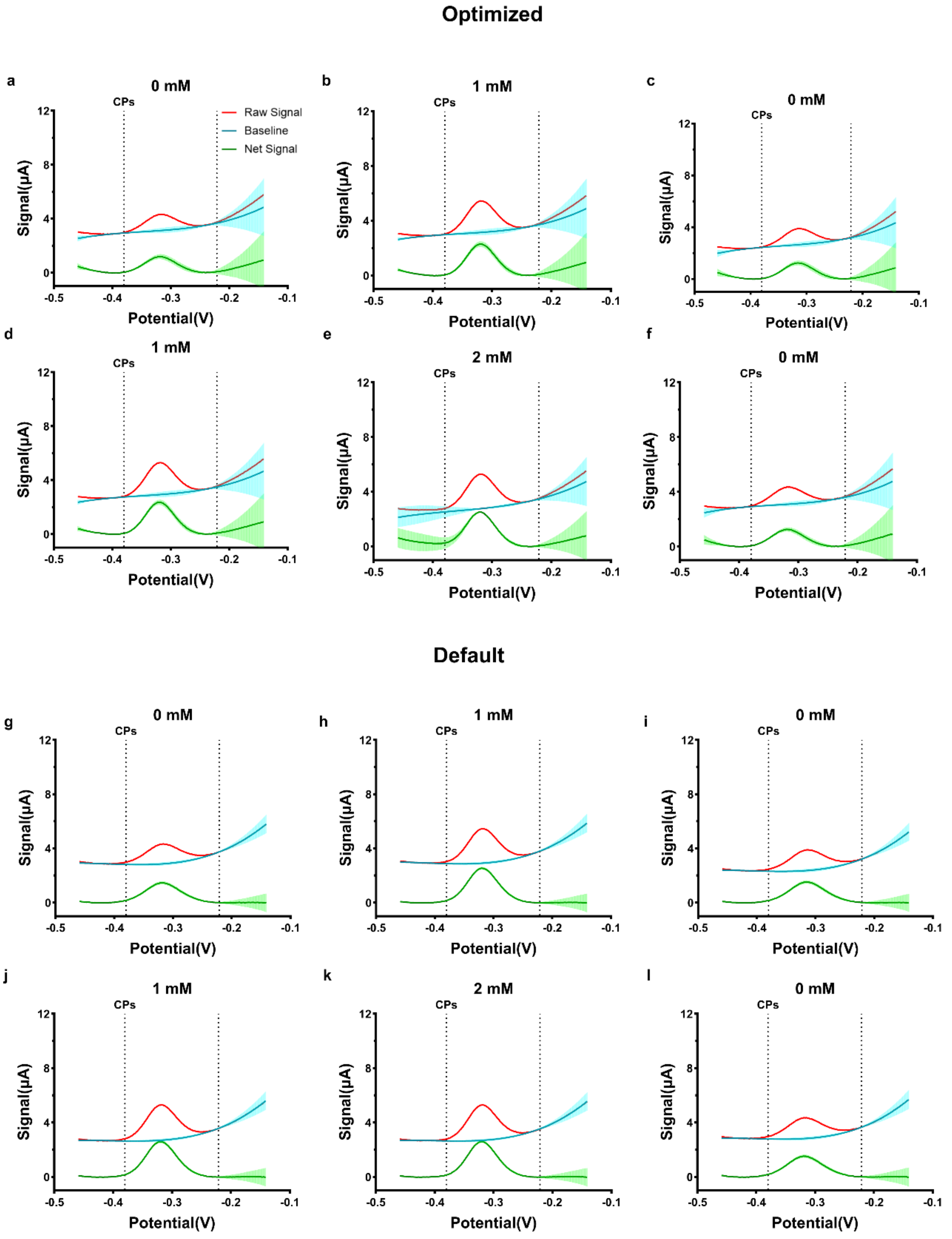

**Figure S20.** *in vivo* day 7 fitting details with default algorithm set and optimized algorithm set. Signals become tilted for all concentrations on day 7.

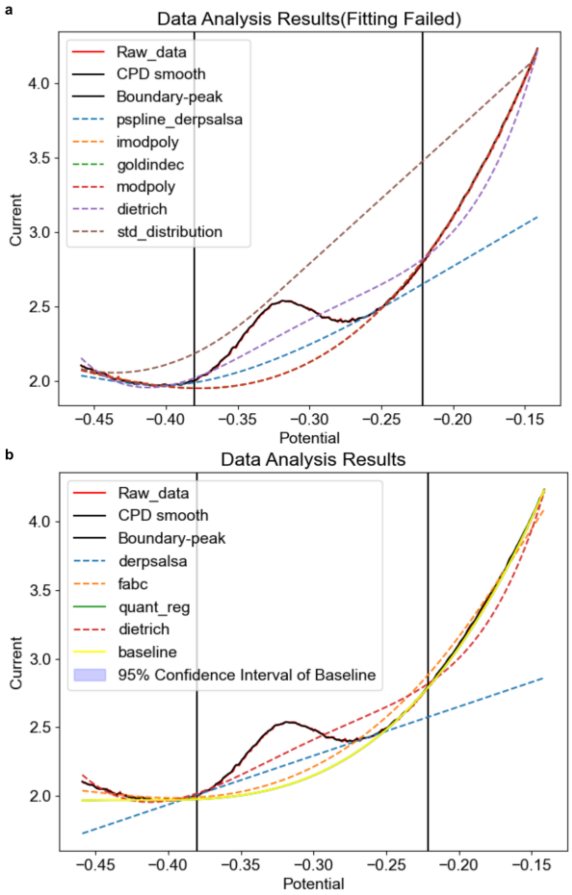

**Figure S21.** Detailed analysis of the fourth data point on day seven. **a** Default algorithm set. In spite of the proper peak region, all baseline fitting algorithms in default algorithm set cannot pass the shape check, which results in a failure in this case. **b** Optimized algorithm set. With the same CPD algorithm and peak region data, one baseline fitting algorithm in the optimal algorithm set passed the shape check and found the peak height.

| Figures | 7a | 7b | S14a | S14b | S14c |
|---|---|---|---|---|---|
| Cosine Similarities | 0.9991 | 0.9638 | 0.9948 | 0.9320 | 0.7822 |
| Figures | S14d | S14e | S14f | S14g | S14h |
| Cosine Similarities | 0.9971 | 0.9993 | 0.9854 | 0.9523 | 0.8765 |

**Table S6.** Cosine similarities over *in vivo* data between A-PACE and MultiTrace. The high cosine similarities show that two tools share the similar analysis trend and the difference of trend is mainly caused by the fluctuation and failure in MultiTrace analysis.

| CPD Algorithm | Baseline Fitting Algorithms |
|---|---|
| BottomUp-rank | derpsalsa, fabc, quant_reg, dietrich, pspline_iarpls |

**Table S7.** Optimized algorithm set for *in vivo* data. BottomUp-rank remains the selected CPD algorithm in both optimal and default algorithms since it finds better CPs. The optimal algorithm set has different baseline fitting algorithms with default algorithm set, which helps it find the proper baselines in the cases where default algorithm set failed.
